# Ciliary signaling-patterned smooth muscle drives tubular elongation

**DOI:** 10.1101/2020.08.31.276295

**Authors:** Ying Yang, Pekka Paivinen, Chang Xie, Alexis Leigh Krup, Tomi P. Makela, Keith E. Mostov, Jeremy F. Reiter

**Affiliations:** Department of Biochemistry and Biophysics, Cardiovascular Research Institute, University of California San Francisco, San Francisco, CA, USA; iCAN Digital Precision Cancer Medicine Flagship, Research Programs Unit, Faculty of Medicine and HiLIFE-Helsinki Institute of Life Science, University of Helsinki, Helsinki, Finland; Department of Anatomy, University of California San Francisco, San Francisco, CA, USA; Chan Zuckerberg Biohub, San Francisco, CA 94158, USA

## Abstract

During development, many tubular organs undergo extensive longitudinal growth to reach their defined length, essential for their function, but how they lengthen is poorly understood. Here, we found that primary cilia are critical for the elongation of the small intestine and esophagus during murine embryonic development. More specifically, HH ligands produced by the epithelium signaled via cilia in the surrounding mesenchyme to pattern the smooth muscle. Like attenuated ciliary HH signaling, partial ablation of the smooth muscle reduced elongation, revealing an essential role for smooth muscle in longitudinal growth. Disruption of cilia, HH signaling or the smooth muscle reduced residual stress within the gut wall, indicating that smooth muscle contributes to the mechanical properties of the developing gut. Reducing residual stress decreased nuclear YAP, an effector of the mechanotransductive Hippo pathway. Removing YAP in the mesenchyme did not affect smooth muscle formation, but attenuated proliferation and elongation, demonstrating that YAP interprets smooth muscle-generated force to promote proliferation. Together, our results reveal that ciliary signaling directs the formation of the smooth muscle layer which, in turn, generates mechanical forces that activate YAP-mediated proliferation. As this interplay of biochemical and mechanical signals drives elongation of both the esophagus and small intestine, we propose that this mechanism may underlie tubular organ elongation generally.

**Highlights:**

- Primary cilia are essential for the elongation of the small intestine and esophagus during embryonic development
- Ciliary signaling patterns the smooth muscle in the developing intestine and esophagus
- The smooth muscle contributes to tissue mechanics
- Smooth muscle-generated strain activates YAP to drive longitudinal growth of the tubular organs

## Introduction

Many of our organs, including the alimentary tract, are epithelial tubes surrounded by mesenchyme. In humans, the endodermal tube post-gastrulation is approximately 1 mm long, and, at birth, the alimentary tract is over 3 m long, representing a >3,000-fold increase in length (FitzSimmons et al., 1988). In contrast, the fetus itself elongates approximately 500-fold in the same time.

In the mouse gut, smooth muscle arises from mesoderm-derived stromal cells beginning at embryonic day (E) 11. During E12-13, these smooth muscle cells organize into a distinct circumferential layer (Chin et al., 2017). Signaling molecules, including Hedgehog (HH) ligands, have been identified to participate in intestinal lengthening (Mao et al., 2010; Walton et al., 2016). Like intestinal elongation, formation of this smooth muscle layer requires HH signaling (Cotton et al., 2017; Huang et al., 2013; Huycke et al., 2019; Mao et al., 2010; Ramalho-Santos et al., 2000). How HH signaling drives both smooth muscle differentiation and intestinal elongation and whether these two events are coupled have been unclear.

Mechanical forces during development can be generated by smooth muscle. During late gestation, smooth muscle generates biophysical cues that instruct the formation of epithelial villi and the second longitudinal smooth muscle layer in the intestine (Huycke et al., 2019; Shyer et al., 2013). Somewhat similarly, smooth muscle is implicated in airway branching (Kim et al., 2015). In the postnatal intestine, applied distractive forces can lead to lengthening (Stark and Dunn, 2012). The sources of mechanical force in development and whether mechanical force contributes to developmental elongation have been unclear.

Mechanical stimuli can activate the Hippo pathway and lead to organ growth (Panciera et al., 2017; Zheng and Pan, 2019). The primary transcriptional effectors of the Hippo pathway are Yes-associated protein (YAP, also called YAP1) and its related co-transcriptional factor TAZ (also called WWTR1) (Huang et al., 2005; Lei et al., 2008). Different mechanical influences cause YAP and TAZ to localize to either the nucleus or the cytoplasm (Dupont et al., 2011; Zhao et al., 2007). Nuclear YAP and TAZ promote cell proliferation and inhibit apoptosis (Dong et al., 2007; Zhao et al., 2008). Although YAP restricts the HH-dependent differentiation of smooth muscle (Cotton et al., 2017), it has been unclear whether, conversely, HH signaling regulates YAP activity.

Vertebrate HH signaling is transduced by a specialized signaling organelle called the primary cilium. Most mammalian cells possess a single, non-motile primary cilium protruding from the cell surface. The primary cilium is comprised of a microtubule-based axoneme built atop a mother centriole and sheathed by a ciliary membrane enriched in specific signaling proteins (Bangs and Anderson, 2017). HH signals trigger the entry of a seven-pass transmembrane protein, Smoothened (SMO), into the cilium where it activates the downstream transcriptional effectors of the HH pathway (Corbit et al., 2005). Although cilia transduce HH signals to pattern the limb and neural tube (García-García et al., 2005; Huangfu and Anderson, 2005; Liem et al., 2012; Liu et al., 2005), whether cilia function in gut development has been unexplored. Therefore, we examined the development of the gut in mouse mutants with defective ciliary signaling.

## Results

### Primary cilia are critical for intestinal elongation during mammalian embryogenesis

Human mutations in *Ciliogenesis associated kinase 1* (*CILK1*, also called *Intestinal cell kinase* or *ICK*) cause lethal endocrine-cerebro-osteodysplasia and short rib polydactyly syndromes (Fu et al., 2019; Lahiry et al., 2009; Oud et al., 2016; Paige Taylor et al., 2016). We and others found that CILK1 is a ciliary kinase important for ciliary morphology and HH signaling (Chaya et al., 2014; Moon et al., 2014; Tong et al., 2017; Yang et al., 2013). To study the effects on intestinal development by impaired cilia, we generated mouse germline and conditional null alleles of *Cilk1* (Figure S1A). Consistent with previous reports (Chaya et al., 2014; Moon et al., 2014), *Cilk1*^-/-^ embryos survived until birth and displayed edema, shortened ribs and polydactyly (Figure S1B). These phenotypes are consistent with developmental roles for CILK1 in regulating ciliogenesis and ciliary HH signaling. In addition to previously described phenotypes, we observed that *Cilk1*^-/-^ embryos exhibited dramatically shortened small intestines (Figure 1A-B). An approximately 50% reduction in intestinal length was completely penetrant in *Cilk1*^-/-^ from E14.5. Other growth parameters, such as intestinal diameter or body length, were unchanged in *Cilk1*^-/-^ embryos (Figure S1C-D), indicating that the intestinal shortening phenotype is not reflective of global growth defects.

**Figure 1.**
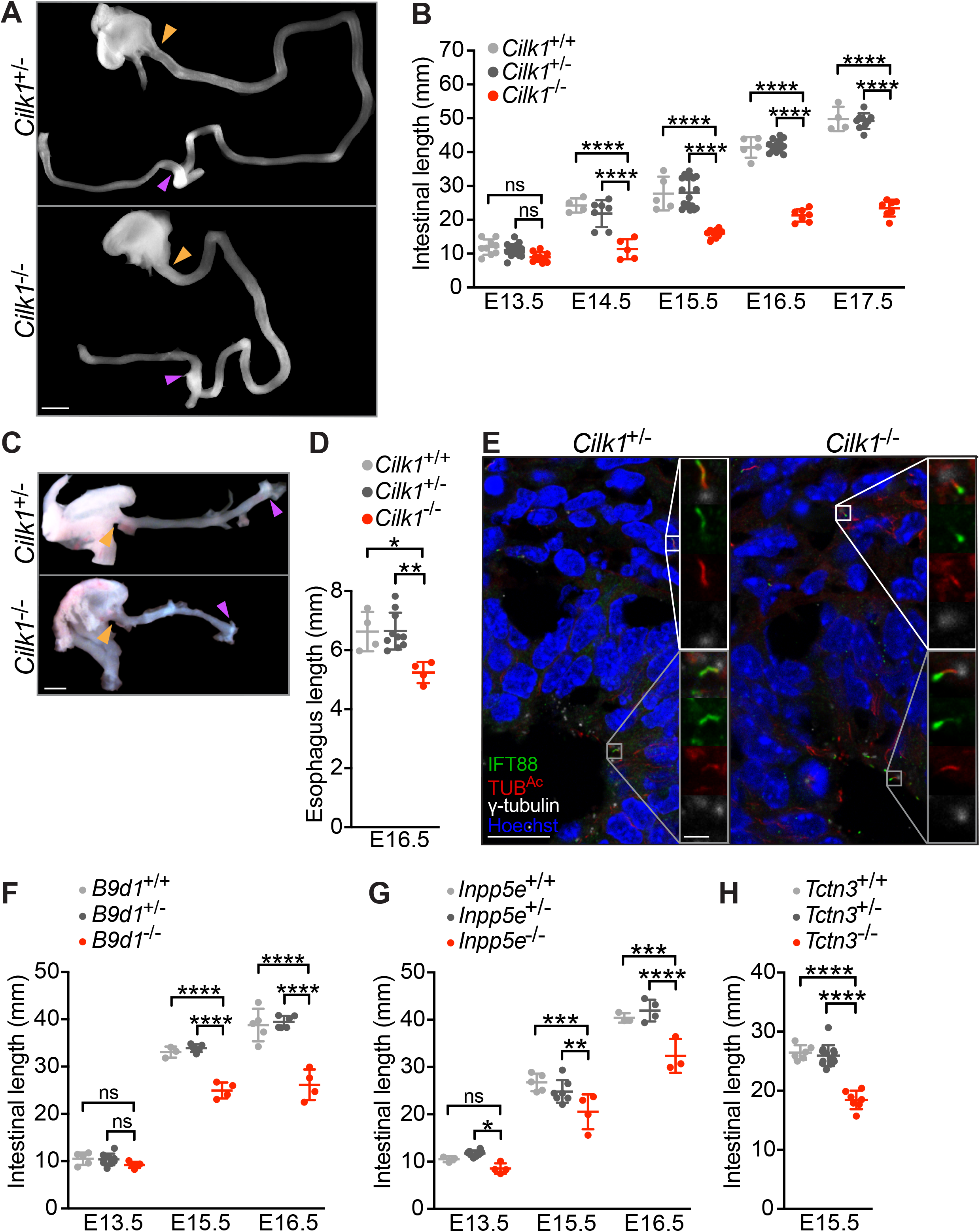
Cilia are essential for the elongation of the small intestine and esophagus in mouse embryos. (A) E14.5 intestines after separation from the mesentery of *Cilk1*^+/-^ and *Cilk1*^-/-^ embryos. The length of the small intestine was measured from the caudal stomach (orange arrowheads) to the rostral cecum (purple arrowheads). Scale bar, 1 mm. (B) Lengths of the small intestines of *Cilk1^+/+^, Cilk1*^+/-^ and *Cilk1*^-/-^ embryos at E13.5, 14.5, 15.5, 16.5 and 17.5. (C) E16.5 esophagi of *Cilk1*^+/-^ and *Cilk1*^-/-^ embryos. The length of the esophagi was measured from the stomach (orange arrowheads) to the pharynx (purple arrowheads). Scale bar, 1 mm. (D) Lengths of the esophagi of *Cilk1^+/+^, Cilk1*^+/-^ and *Cilk1*^-/-^ embryos at E16.5. (E) Immunofluorescence staining of E13.5 intestinal cross sections from *Cilk1*^+/-^ and *Cilk1*^-/-^ embryos for IFT88 (green), ciliary axonemes (TUB^Ac^, red), basal bodies (γ-tubulin, white) and nuclei (Hoechst, blue). Right, higher magnifications of boxed cilia. White boxes highlight mesenchymal cilia and grey boxes highlight epithelial cilia. Scale bar of larger images, 5 μm. Scale bar of enlarged images, 2 μm. (F) Lengths of the small intestines of *B9d1*^+/+^, *B9d1*^+/-^ and *B9d1*^-/-^ embryos at E13.5, 15.5 and 16.5. (G) Lengths of the small intestines of *Inpp5e^+/+^, Inpp5e*^+/-^ and *Inpp5e*^-/-^ embryos at E13.5, 15.5 and 16.5. (H) Lengths of the small intestines of *Tctn3^+/+^, Tctn3*^+/-^ and *Tctn3*^-/-^ embryos at E15.5. (B), (D), (F), (G) and (H), each point represents the intestinal or esophageal length of one embryo. Horizontal bars indicate means ± SD. ns, p > 0.05; *p < 0.05; **p < 0.01; *** p < 0.001; **** p < 0.0001 by one-way ANOVA Tukey’s multiple comparisons test performed for each embryonic stage.

To investigate another part of the gut, we examined the developing esophagus. Like the small intestines, *Cilk1*^-/-^ esophagi were shorter than those of control (Figure 1C-D). Thus, CILK1 promotes elongation in multiple parts of the developing gut.

To begin to investigate how CILK1 regulates intestinal elongation, we examined the intestinal cilia in *Cilk1* mutants to assess whether CILK1 regulates the composition of cilia in the developing intestine. A ciliary protein, Intraflagellar Transport 88 (IFT88), was distributed along the length of control cilia. In contrast, IFT88 abnormally accumulated in the ciliary tips in both *Cilk1*^-/-^ intestine epithelial (Figure 1E, grey box) and mesenchymal cells (Figure 1E, white box). This altered distribution of IFT88 indicates that CILK1 controls ciliary composition in the developing gut. As IFT88 is critical for intraciliary trafficking and ciliary signaling transduction, these results further raise the possibility that the requirement for of CILK1 in tubular elongation could be dependent on cilia.

To test whether the role of CILK1 in intestinal development reflects its ciliary function, we examined intestines of other mutants affecting ciliary function, including mouse embryos lacking *B9 domain-containing 1* (*B9d1*) (Dowdle et al., 2011), or *Inositol polyphosphate 5-phosphatase* (*Inpp5e*) (Jacoby et al., 2009). In addition, we generated a loss-of-function allele of *Tectonic 3* (*Tctn3*) (Figure S2A), encoding a component of the ciliary transition zone we previously identified (Garcia-Gonzalo et al., 2011). Consistent with a previous report (Wang et al., 2018), the *Tctn3*^-/-^ embryos displayed microphthalmia and polydactyly at E13.5 (Figure S2B).

We selected the *B9d1, Inpp5e* and *Tctn3* mouse mutants because the homozygous null mutants survive to gestational ages at which intestinal length can be ascertained and because they encode proteins that participate in ciliary signaling through distinct mechanisms. INPP5E generates a cilium-enriched phosphoinositide important for ciliary signaling (Chávez et al., 2015; Dyson et al., 2017; Garcia-Gonzalo et al., 2015). B9D1 and TCTN3 are components of the ciliary transition zone controlling ciliary composition (Garcia-Gonzalo et al., 2011). Consistent with a function of TCTN3 at the transition zone, immunofluorescence staining for ARL13B, a transition zone-dependent ciliary component, revealed that E13.5 *Tctn3*^-/-^ embryos displayed reduced numbers of ARL13B-positive cilia (Figure S2C).

Like *Cilk1* null mutants, *B9d1, Inpp5e* and *Tctn3* mutants all exhibited shortened small intestines (Figure 1F-H). At E15.5, *Cilk1* mutant intestines were shortened by 43%, whereas *B9d1* mutant intestines were shortened by 26%, *Inpp5e* mutant intestines were shortened by 17% and *Tctn3* mutant intestines were shortened by 29%. Thus, disrupting multiple aspects of ciliary function all resulted in shortened intestines, indicating that cilia promote elongation.

### Mesenchymal cilia are essential for HH signal transduction and tubular elongation

Mesenchymal cells, which are the main HH responsive cells in the developing gut (Madison et al., 2005), possessed cilia throughout development (Figure S3, white boxes). In contrast, epithelial cells, which produce the HH ligands (Motoyama et al., 1998), possessed cilia at E13.5 (Figure 1E) but not at E18.5 (Figure S3, grey boxes), consistent with a previous study (Komarova and Vorob’ev, 1995). As cilia were present in both the endoderm-derived epithelium and the mesoderm-derived mesenchyme surrounding the epithelium during early intestinal development, roles for cilia in intestinal elongation could reflect functions in the epithelium, the mesenchyme or both compartments. To distinguish between these possibilities, we removed CILK1 from either the epithelium or the mesenchyme and assessed intestinal length.

To remove CILK1 specifically from the epithelium, we generated *Shh*^Cre^ *Cilk1*^lox/lox^ embryos, in which *Shh*^Cre^ (Harfe et al., 2004) deleted *Cilk1* specifically in the endodermal epithelium (Figure S4A). Consistent with paracrine HH signaling in the intestine (Madison et al., 2005), the expression of HH target genes *Gli1* and *Ptch1* was enriched in the stromal compartment (Figure S4A). *Gli1* and *Ptch1* expression levels were unchanged in *Shh*^Cre^ *Cilk1*^lox/lox^ embryos, and abrogating *Cilk1* expression in the intestinal epithelium did not alter the length of *Shh*^Cre^ *Cilk1*^lox/lox^ intestines at E18.5 (Figure S4A-B), suggesting that CILK1 in the epithelium is dispensable for intestinal elongation.

To remove CILK1 specifically from the mesenchyme, we generated *Dermo1*^Cre^ *Cilk1*^lox/lox^ embryos. *Dermo1*^Cre^ (Yu et al., 2003) was active in mesoderm-derived intestinal mesodermal cells, as revealed by Cre-dependent tdTomato expression (Figure S5A). In *Dermo1*^Cre^ *Cilk1*^lox/lox^ intestine, the expression of *Cilk1* was reduced as were the HH target genes *Gli1* and *Ptch1* (Figure 2A), indicating that CILK1 functions in the mesenchyme to transduce HH signals. Consistent with the essential role of cilia in HH signaling in other contexts, these results suggest that mesenchymal cilia interpret HH signals in the developing gut.

**Figure 2.**
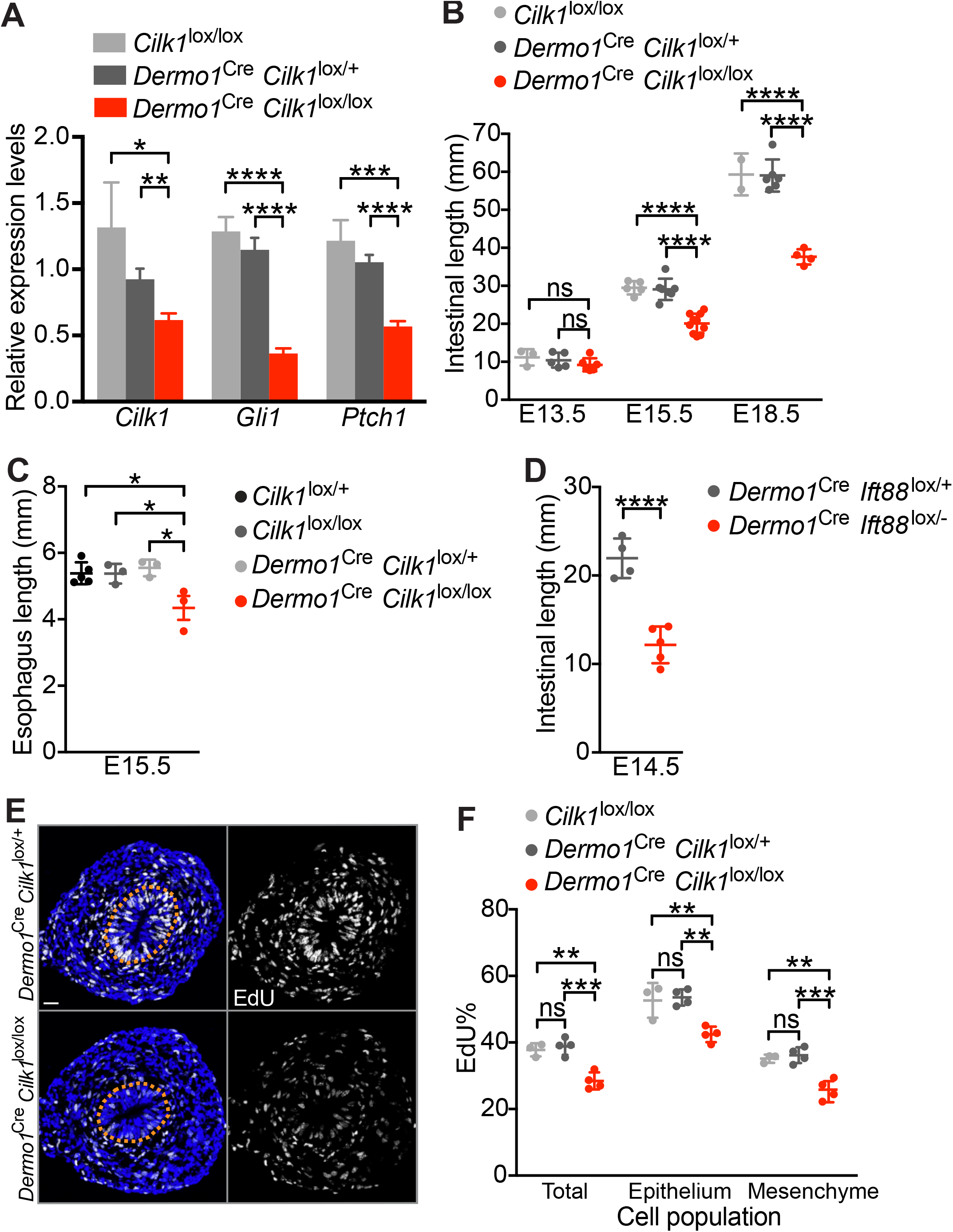
CILK is specifically required in the mesenchymal cells to promote Hh signaling responsiveness, proliferation and intestinal elongation. (A) Relative levels of *Cilk1*, *Gli1* and *Ptch1* mRNA in E15.5 control (*Cilk1*^lox/lox^, n=4 and *Dermo1*^Cre^ *Cilk1*^lox/+^, n=6) and *Dermo1*^Cre^ *Cilk1*^lox/lox^ (n=8) intestines. Values are presented as means ± SEM. p<0.0001 by two-way ANOVA test with genotype as a source of variation. *p < 0.05; **p < 0.01 and ****p<0.0001 by unpaired t test. (B) Lengths of E13.5, E15.5 and E18.5 control (*Cilk1*^lox/lox^ and *Dermo1*^Cre^ *Cilk1*^lox/+^) and *Dermo1*^Cre^ *Cilk1*^lox/lox^ small intestines. (C) Lengths of E15.5 control (*Cilk1*^lox/+^, *Cilk1*^lox/lox^ and *Dermo1*^Cre^ *Cilk1*^lox/+^) and *Dermo1*^Cre^ *Cilk1*^lox/lox^ esophagi. (D) Lengths of E14.5 *Dermo1*^Cre^ *Ift88*^lox/+^ and *Dermo1*^Cre^ *Ift88*^lox/-^ small intestines. **** p < 0.0001 by unpaired t test. (E) Immunofluorescence staining for EdU (white) and nuclei (Hoechst, blue) in E13.5 *Dermo1*^Cre^ *Cilk1*^lox/+^ and *Dermo1*^Cre^ *Cilk1*^lox/lox^ intestines. Scale bar, 25 μm. Orange dotted lines outline the epithelium. (F) The percentage of EdU-positive cells in the whole intestine, the epithelial cells and the mesenchymal cells in *Cilk1*^lox/lox^ (n=3), *Dermo1*^Cre^ *Cilk1*^lox/+^ (n=4) and *Dermo1*^Cre^ *Cilk1*^lox/lox^ (n=4) embryos at E13.5. For each intestine, we averaged the percentages from three sections. (B), (C), (D) and (F) Each point in the scatter plots represents the value from an individual embryonic intestine or espohagus. Horizontal bars indicate means ± SD. ns, p > 0.05; *p < 0.05; **p < 0.01; *** p < 0.001 and ****p<0.0001 by ordinary oneway ANOVA Tukey’s multiple comparisons test for each embryonic stage or individual cell population.

Elongation of intestines in *Shh*^Cre^ *Cilk1*^lox/lox^ embryos was comparable to controls (Figure S4B). Unlike *Shh*^Cre^ *Cilk1*^lox/lox^ embryos, but like *Cilk1*^-/-^ embryos (Figure 1A-B), *Dermo1*^Cre^ *Cilk1*^lox/lox^ embryos exhibited shortened intestines (Figure 2B). At E15.5, *Dermo1*^Cre^ *Cilk1*^lox/lox^ intestines were shortened by 31% (Figure 2B) and *Cilk1*^-/-^ intestines were shortened by 43% (Figure 1B). Thus, intestinal lengthening during development depends upon CILK1 in the mesoderm-derived mesenchyme.

As in the intestine, depletion of CILK1 in the mesoderm-derived cells in *Dermo1*^Cre^ *Cilk1*^lox/lox^ embryos shortened esophagi at E15.5 (Figure 2C). Thus, the lengthening of esophagus during development also depends upon ciliary signaling in the mesenchyme.

As CILK1 is required in the mesenchyme for both HH signal transduction and elongation, we hypothesized that ciliary HH signaling in the mesenchyme is essential for intestinal elongation. To test this hypothesis, we removed IFT88, essential for cilia assembly and maintenance (Pazour et al., 2000), in the mesenchyme (by generating *Dermo1*^Cre^ *Ift88*^lox/-^ embryos). *Dermo1*^Cre^ *Ift88*^lox/-^ embryos, like *Dermo1*^Cre^ *Cilk1*^lox/lox^ embryos, exhibited shortened intestines at E14.5 (Figure 2D). Together, these results indicate that mesenchymal cilia transduce HH signals to drive elongation.

How might mesenchymal ciliary signaling promote elongation? Organ size depends on both proliferation and cell survival. Programmed cell death plays a role in regulating organ size (Hakem et al., 1998; Haydar et al., 1999; Ishizuya-Oka and Ueda, 1996; Kuida et al., 1998; Porteous et al., 2000). To assess whether increased apoptosis could account for decreased intestinal elongation, we quantified the prevalence of apoptotic cells marked by cleaved Caspase-3. We detected no increased in apoptosis in *Dermo1*^Cre^ *Cilk1*^lox/lox^ guts (Figure S6).

In addition to cell death, organ growth reflects cell proliferation rates. To examine whether cilium-dependent intestinal elongation is driven by proliferation, we quantified proliferating cells using 5-ethynyl-2’-deoxyuridine (EdU) labeling of *Dermo1*^Cre^ *Cilk1*^lox/lox^ and littermate control embryos at E13.5, prior to the detection of a difference in intestinal length. The proportion of EdU-labelled cells was reduced in *Dermo1*^Cre^ *Cilk1*^lox/lox^ intestines (Figure 2E-F), indicating that CILK1 promotes intestinal cell proliferation. Interestingly, the decreased proliferation in *Dermo1*^Cre^ *Cilk1*^lox/lox^ embryos was not restricted to the mesenchymal cells where CILK1 was depleted, but was also observed in the epithelial cells (Figure 2E-F). These results demonstrate that mesenchymal CILK1 is essential for proliferation of both the mesenchyme and the underlying epithelium in the developing intestine, suggesting that elongation depends on cilium-promoted proliferation.

### Ciliary signaling patterns the smooth muscle

As HH signaling has been shown to regulate the patterning of smooth muscle cells (Cotton et al., 2017; Huang et al., 2013; Huycke et al., 2019; Mao et al., 2010; Ramalho-Santos et al., 2000), we explored whether cilia participate in intestinal smooth muscle development. We stained for α-smooth muscle actin (αSMA), a marker of smooth muscle, at E13.5, prior to altered gut elongation in cilia mutants. We found that smooth muscle organization was disrupted in *Cilk1*^-/-^ gut, as measured by increased variance of tangential alignment and increased radial distribution (Figure 3A-B). This patterning defect was not due to the presence of more smooth muscle cells (Figure 3C), but altered distribution in the gut wall (Figure 3B). Therefore, CILK1 is not essential for smooth muscle differentiation or survival, but rather for its patterning and spatial organization.

**Figure 3.**
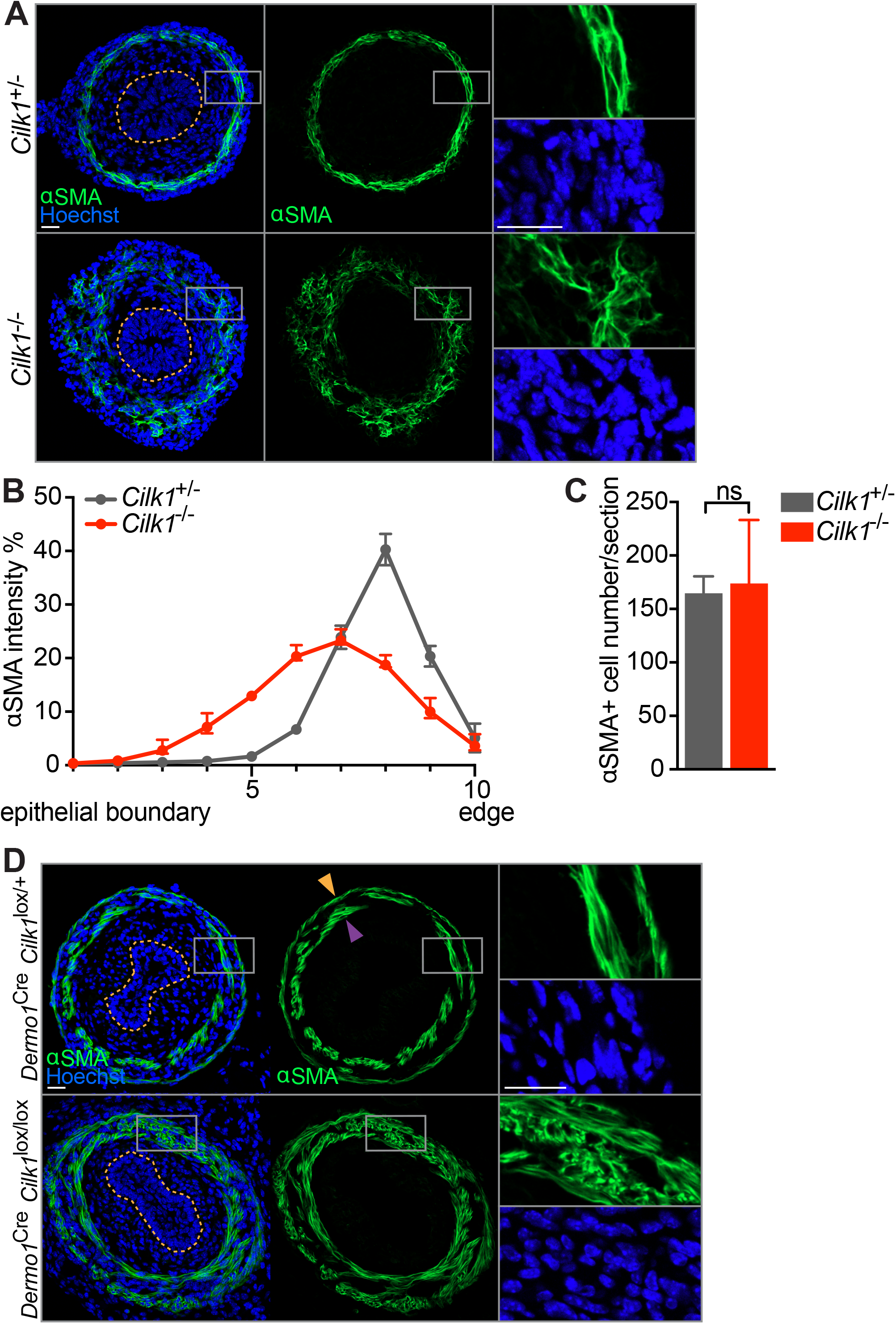
The circumferential smooth muscle cell layer is disrupted in the cilia-deficient intestines at E13.5. (A) Immunofluorescence staining for smooth muscle (αSMA, green) and nuclei (Hoechst, blue) in E13.5 *Cilk1*^+/-^ and *Cilk1*^-/-^ intestines. (B) The distribution of αSMA relative staining intensity from the epithelial boundary to the outer section edge of E13.5 *Cilk1*^+/-^ (n=4) and *Cilk1*^-/-^ (n=4) intestines, subdivided into deciles. For each intestine, the αSMA intensity fraction in each decile was averaged from 3-8 sections. Values are presented as means ± SD. X axis, decile number. (C) The number of αSMA-positive cells per section from E13.5 *Cilk1*^+/-^ (n=3) and *Cilk1*^-/-^ (n=3) intestines. For each embryonic intestine, the number of αSMA-positive cells was averaged from 3-8 stained sections. Values are presented as means ± SD. ns, p > 0.05 by unpaired t test. (D) Immunofluorescence staining of E15.5 *Dermo1*^Cre^ *Cilk1*^lox/+^ and *Dermo1*^Cre^ *Cilk1*^lox/lox^ esophagi for smooth muscle (αSMA, green) and nuclei (Hoechst, blue). Upper panel, arrowheads indicate two layers of esophageal muscularis: lamina muscularis interna (purple) and the lamina muscularis externa (orange). (A) and (D), right, higher magnifications of boxed regions. Orange dotted lines indicate the epithelial boundary. Scale bars, 25 μm.

To test whether the role of CILK1 in smooth muscle development is shared by other cilia-related proteins, we examined smooth muscle in *Tctn3*^-/-^ gut. As with CILK1, TCTN3 was required for organization of the circumferential smooth muscle (Figure S7A). Together, these results indicate that cilia are required for the formation of the circumferential smooth muscle in the developing gut.

To assess whether mesenchymal cilia are specifically required for smooth muscle development, as they are for elongation, we examined smooth muscle in E13.5 in control and *Dermo1*^Cre^ *Cilk1*^lox/lox^ guts (Figure S7B). Similar to *Cilk1*^-/-^ smooth muscle, the smooth muscle of *Dermo1*^Cre^ *Cilk1*^lox/lox^ guts was disorganized with increased radial distribution and disrupted circumferential alignment, indicating that cilia functionally specifically in the mesenchyme to pattern smooth muscle.

To examine whether mesenchymal ciliary signaling also patterns esophageal smooth muscle, we stained for αSMA in esophagi. As reported before (Huycke et al., 2019; Kablar et al., 2000), at E15.5, αSMA was predominantly expressed by two layers of esophageal smooth muscle (Figure 3D upper panel): lamina muscularis interna (purple arrowhead) and lamina muscularis externa (orange arrowhead). Both layers of lamina muscularis surround the esophageal epithelium (Figure 3D upper panel). In *Cilk1*^-/-^ esophagus, the spatial separation of the two smooth muscle layers was decreased (Figure 3D lower panel). In addition, some smooth muscle fibers were misoriented in *Cilk1*^-/-^ esophagus (Figure 3D lower panel). Thus, as in the intestine, CILK1 regulates the spatial organization of smooth muscle in the developing esophagus.

### Mesenchymal HH signaling is required for development of smooth muscle

The essential role for cilia and CILK1 in transducing HH cues raised the possibility that HH signaling in the mesenchyme patterns the smooth muscle and elongates the gut. Previous work demonstrated that depletion of epithelial HH ligands shortens intestinal length (Mao et al., 2010). To test whether HH signals transduced by mesenchymal cilia promote elongation, we removed Smoothened (SMO), an essential component of the HH signal transduction pathway, from the mesenchyme. After generating *Dermo1*^Cre^ *Smo*^lox/lox^ embryos and assessing their intestinal length at E14.5, we found that these intestines were shortened by approximately 55% (Figure 4A). Thus, mesenchymal HH signaling promotes gut elongation.

**Figure 4.**
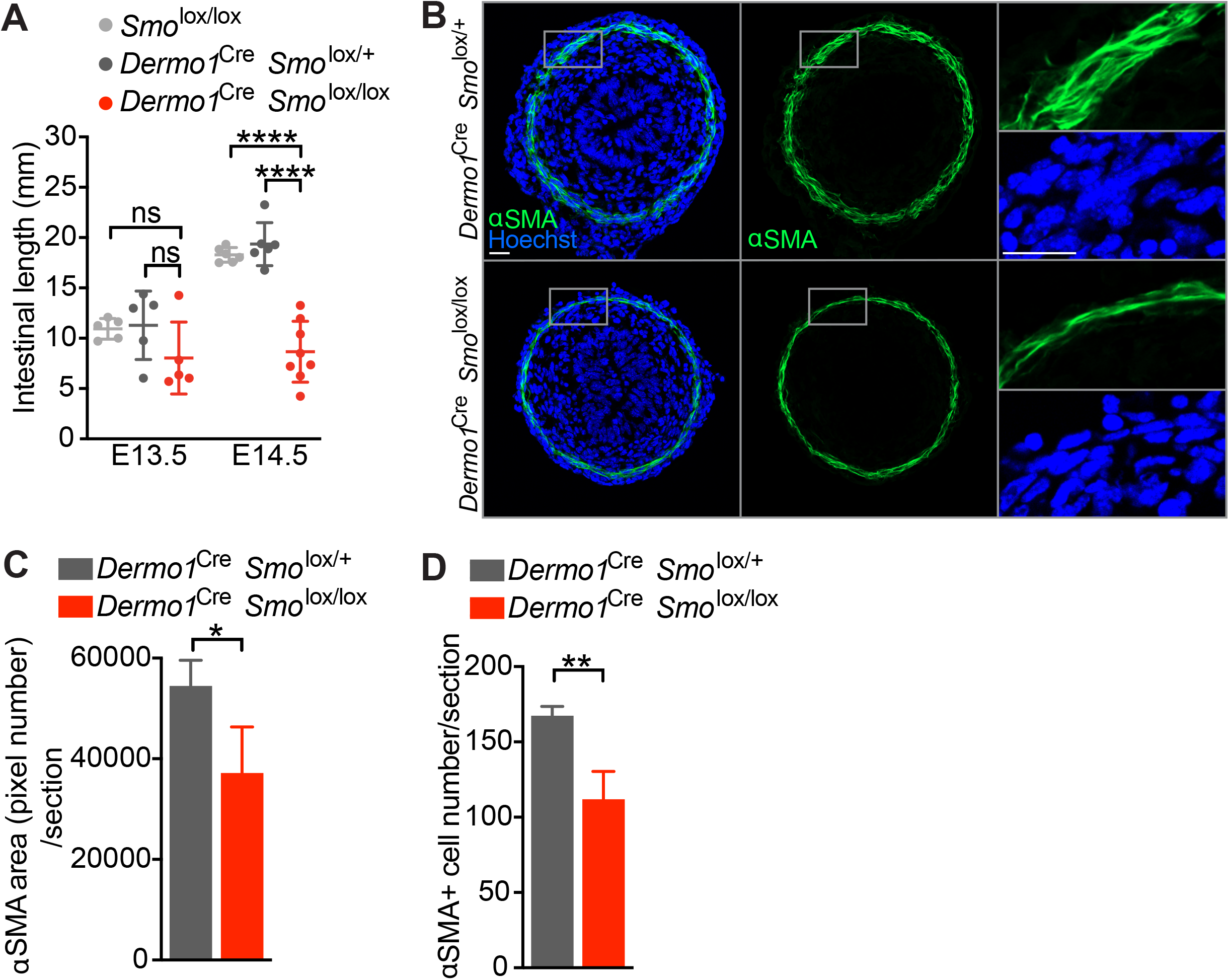
SMO acts in the mesenchyme to promote intestinal elongation and the formation of the circumferential smooth muscle. (A) Lengths of E13.5 and E14.5 control (*Smo*^lox/lox^ and *Dermo1*^Cre^ *Smo*^lox/+^) and *Dermo1*^Cre^ *Smo*^lox/lox^ small intestines. Each point in the scatter plots represents the value from an individual embryo. Horizontal bars indicate means ± SD. ns, p > 0.05; **** p < 0.0001 by one-way ANOVA Tukey’s multiple comparisons test performed for each embryonic stage. (B) Immunofluorescence staining for smooth muscle (αSMA, green) and nuclei (Hoechst, blue) in E13.5 *Dermo1*^Cre^ *Smo*^lox/+^ and *Dermo1*^Cre^ *Smo*^lox/lox^ intestines. Right, higher magnifications of boxed regions. Scale bars, 25 μm. (C) The area of αSMA staining per section from E13.5 *Dermo1*^Cre^ *Smo*^lox/+^ (n=3) and *Dermo1*^Cre^ *Smo*^lox/lox^ (n=3) intestines. For each embryonic intestine, the αSMA area (pixel number) was averaged from 5-8 sections. (D) The number of αSMA-positive cells per section from E13.5 control (*Dermo1*^Cre^ *Smo*^lox/+^, n=3) and *Dermo1*^Cre^ *Smo*^lox/lox^ (n=3) intestines. For each intestine, the number of αSMA-positive cells was averaged from 5-8 sections. (C-D) Values are presented as means ± SD. * p < 0.05; **p < 0.01 by unpaired t test.

To test whether disrupting SMO also affects smooth muscle development, we analyzed smooth muscle in *Dermo1*^Cre^ *Smo*^lox/lox^ and control embryos at E13.5. In *Dermo1*^Cre^ *Smo*^lox/lox^ embryos, the circumferential smooth muscle layer was thinner (Figure 4B-C) and the number of αSMA-expressing cells was reduced (Figure 4D). Thus, both cilia and HH signaling are essential to pattern the intestinal circumferential smooth muscle.

### Elongation depends on smooth muscle integrity

Do disruptions in mesenchymal cilia or HH signaling independently impair smooth muscle development and decrease elongation, or do the defects in smooth muscle cause decreased elongation? To distinguish between these two possibilities, we tested whether smooth muscle itself is critical for elongation. Specifically, we partially ablated smooth muscle by expressing attenuated diphtheria toxin (DTA176) in smooth muscle under the control of the *Myosin heavy polypeptide 11* (*Myh11*) regulatory elements (*Myh11*^Cre-EGFP^ *R26*^DTA176/+^ embryos). To validate decreased smooth muscle, we co-stained the E15.5 intestine for αSMA and GFP. In the *Myh11*^Cre-EGFP^ intestine, most αSMA-expressing cells also expressed GFP, indicating the *Myh11*^Cre-EGFP^ is active in most smooth muscle cells (Figure 5A, upper panel). As expected, GFP expressing cells were mostly absent in *Myh11*^Cre-EGFP^ *R26*^DTA176/+^ embryos (Figure 5A, lower panel). The number of αSMA-positive cells in *Myh11*^Cre-EGFP^ *R26*^DTA176/+^ embryos was reduced to 63% of controls (Figure 5B).

**Figure 5.**
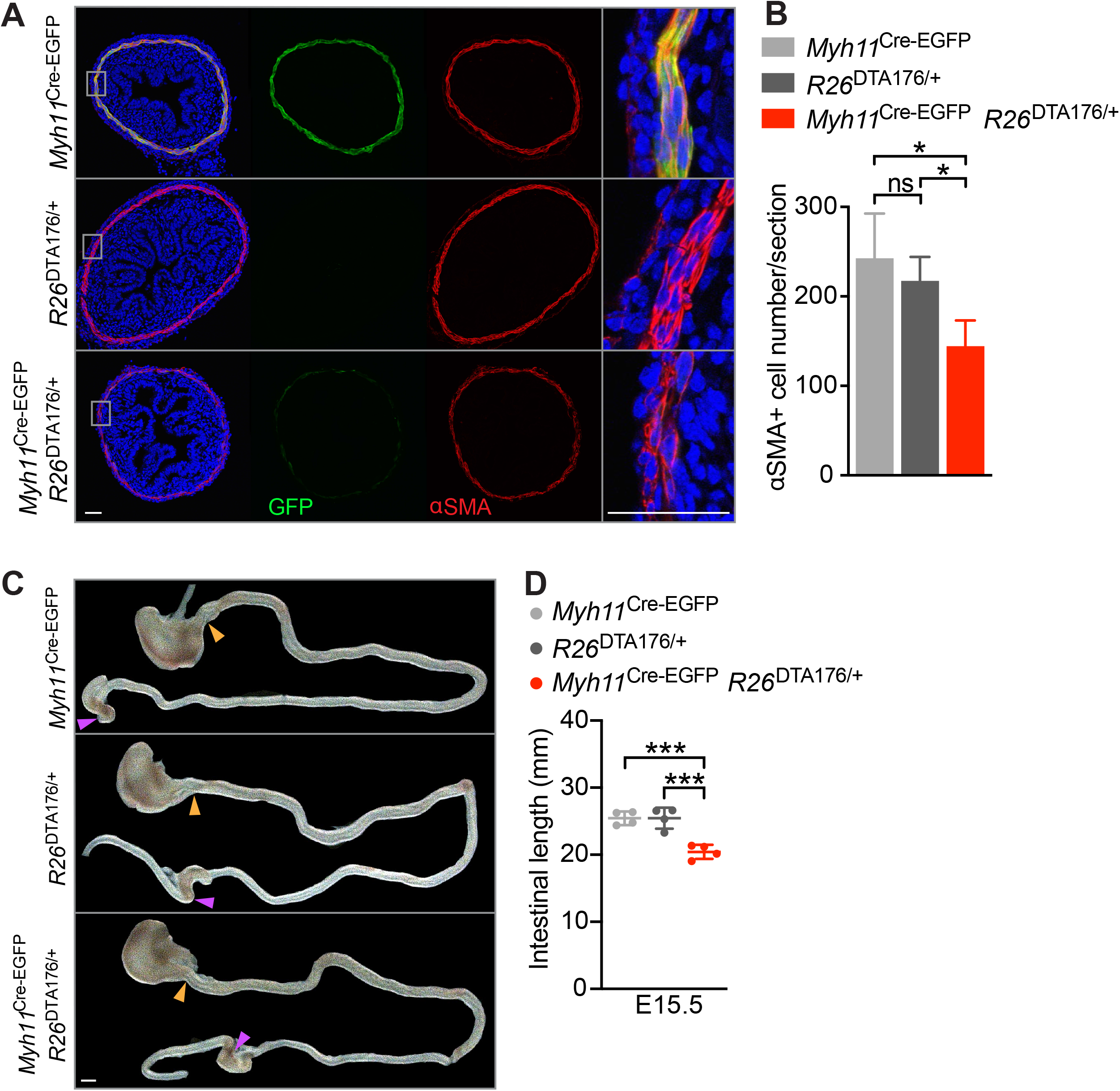
Smooth muscle is required for intestinal elongation. (A) Immunofluorescence staining of E15.5 control (*Myh11*^Cre-EGFP^ and *R26*^DTA176/+^) and *Myh11*^Cre-EGFP^ *R26*^DTA176/+^ intestines for GFP (green), smooth muscle (αSMA, red) and nuclei (Hoechst, blue). Magnifications of the boxed region shown at right. Scale bars, 50 μm. (B) The number of αSMA-positive cells per E15.5 control (*Myh11*^Cre-EGFP^, n=3 and *R26*^DTA176/+^, n=3) and *Myh11*^Cre-EGFP^ *R26*^DTA176/+^ (n=3) intestinal section. For each intestine, the number of αSMA-positive cells was averaged from 2-4 sections. Values are presented as means ± SD. The p value by ordinary one-way ANOVA is 0.0398. ns p > 0.05, * p < 0.05 by unpaired t test. (C) Intestines from E15.5 control (*Myh11*^Cre-EGFP^ and *R26*^DTA176/+^) and *Myh11*^Cre-EGFP^ *R26*^DTA176/+^ embryos. Scale bar, 1 mm. The length of the small intestine was measured from the end of the stomach (orange arrowheads) to the beginning of cecum (purple arrowheads). (D) Lengths of E15.5 control (*Myh11*^Cre-EGFP^ and *R26*^DTA176/+^) and *Myh11*^Cre-EGFP^ *R26*^DTA176/+^ small intestines. Each point in the scatter plots represents the value from an individual embryo. Values are presented as means ± SD. *** p < 0.001 by ordinary one-way ANOVA Tukey’s multiple comparisons test.

Partially ablating the smooth muscle of *Myh11*^Cre-EGFP^ *R26*^DTA176/+^ embryos did not grossly alter embryonic development at E15.5 (Figure S8A). However, *Myh11*^Cre-EGFP^ *R26*^DTA176/+^ embryos exhibited shortened small intestines (Figure 5C-D). Thus, the smooth muscle is critical for elongation, indicating that ciliary HH signaling patterns the smooth muscle to drive elongation.

### Cilia- and HH-dependent patterning of the smooth muscle determines residual stress

To understand how smooth muscle promotes elongation, we explored how the circumferential smooth muscle contributes to mechanical characteristics. Like other muscles, both contraction and passive biophysical properties define the forces produced by smooth muscle (Kobelev et al., 2011). In cultured embryonic chicken intestine, the contractions of the circumferential smooth muscle promote anisotropic growth (Khalipina et al., 2019). To assess mouse smooth muscle contractility, we stained for phosphorylated Myosin Light Chain (phospho-MLC), a component of activated myosin (Allen and Walsh, 1994). Smooth muscle was positive for phospho-MLC at E18.5, but negative at E13.5 (Figure S9), suggesting that the circumferential smooth muscle promotes intestinal elongation before it becomes contractile.

In addition to active contractions, passive cellular and endomysial elasticity contribute to muscle mechanical properties. Tubular organs, including the gut, exhibit a force called residual stress even in the absence of external loads (Gregersen and Kassab, 1996). We hypothesized that the smooth muscle is a source of circumferential residual stress. To test this hypothesis, we measured the opening angles (θ) resulting from longitudinally opening gut segments (Figure 6A), indicators of tensile residual stress (Fung, 1991; Han and Fung, 1991; Zhao et al., 2002).

**Figure 6.**
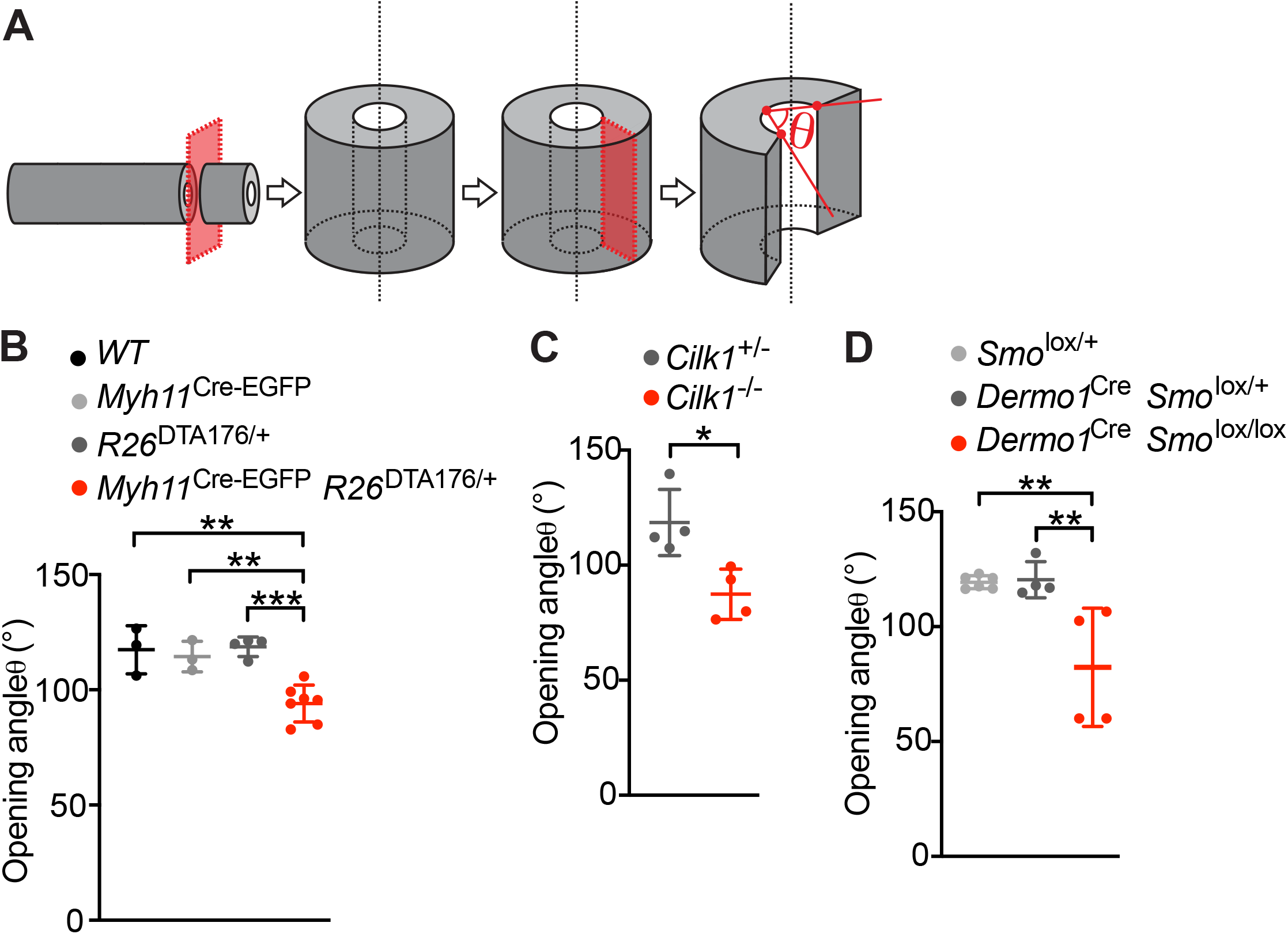
CILK1 function, HH signaling and smooth muscle all contribute to intestinal residual stress. (A) Schematic of method for measuring the opening angle to assess residual stress. We isolated approximately 1mm long sections of E13.5 small intestine, cut the intestinal wall axially and measured the opening angle (θ), with the vertex at the midpoint of the inner epithelial surface and the rays intersecting the innermost edges of the cut wall. (B) The opening angles of intestinal segments from E13.5 control (wild type, n=3, *Myh11*^Cre-EGFP^, n=3 and *R26*^DTA176/WT^, n=4) and *Myh11*^Cre-EGFP^ *R26*^DTA176/+^ (n=7) embryos. (C) The opening angles of intestinal segments from E13.5 *Cilk1*^+/-^ (n=4) and *Cilk1*^-/-^ (n=4) embryos. (D) The opening angles of intestinal segments from E13.5 (*Smo*^lox/+^, n=6 and *Dermo1*^Cre^ *Smo*^lox/+^, n=4) and *Dermo1*^Cre^ *Smo*^lox/lox^ (n=4) embryos. (B-D) For each intestine, we calculated the median angle of 5-15 segments. Values are presented as means ± SD. * p < 0.05; ** p < 0.01; *** p < 0.001 by ordinary one-way ANOVA Sidak’s multiple comparisons test, except in (C) * p < 0.05 by unpaired t test.

To assay whether intestinal residual stress is produced by smooth muscle, we measured the opening angle of intestinal segments of control and *Myh11*^Cre-EGFP^ *R26*^DTA176/+^ embryos. The opening angles were reduced in *Myh11*^Cre-EGFP^ *R26*^DTA176/+^ embryos (94.1° ± 8.0) as compared to controls (wild type 117.4° ± 10.4, *Myh11*^Cre^ 114.4° ± 6.6, *R26*^DTA176/+^ 118.6° ± 4.2) (Figure 6B). Therefore, smooth muscle contributes to circumferential residual stress.

As partial ablation of the smooth muscle decreased residual stress in the intestine, we predicted that altered cilia function or mesenchymal HH signaling, both of which disrupt smooth muscle patterning, should also decrease residual stress. To begin to test this prediction, we examined the opening angles of *Cilk1*^-/-^ intestines and found that compromising cilia function by loss of CILK1 reduced the opening angle by 31.1° ± 9.0 (Figure 6C).

We also examined the opening angle of *Dermo1*^Cre^ *Smo*^lox/lox^ intestines and found that disruption of mesenchymal HH signaling also reduced intestinal residual stress (Figure 6D). Therefore, disruption of the smooth muscle by genetically ablation, impaired cilia, or HH signaling, each reduces intestinal residual stress and compromises elongation. Together, these data raised the interesting possibility that residual stress produced by the circumferential smooth muscle promotes longitudinal growth.

### Smooth muscle activates YAP in the gut

In Drosophila development, mechanical strain is interpreted by the Hippo pathway (Fletcher et al., 2018). The primary effector of the Hippo pathway is the transcriptional co-activator YAP, which translocates to the nucleus upon activation. To assess whether the Hippo pathway also interprets mechanical stresses in the developing gut, we stained E13.5 intestine for YAP. Interestingly, the nuclear localization of YAP depended on radial location: inner mesenchymal cells between the epithelium and the smooth muscle displayed higher levels of nuclear YAP, and more peripheral smooth muscle cells possessed lower levels of nuclear YAP (Figure 7A, top panel).

**Figure 7.**
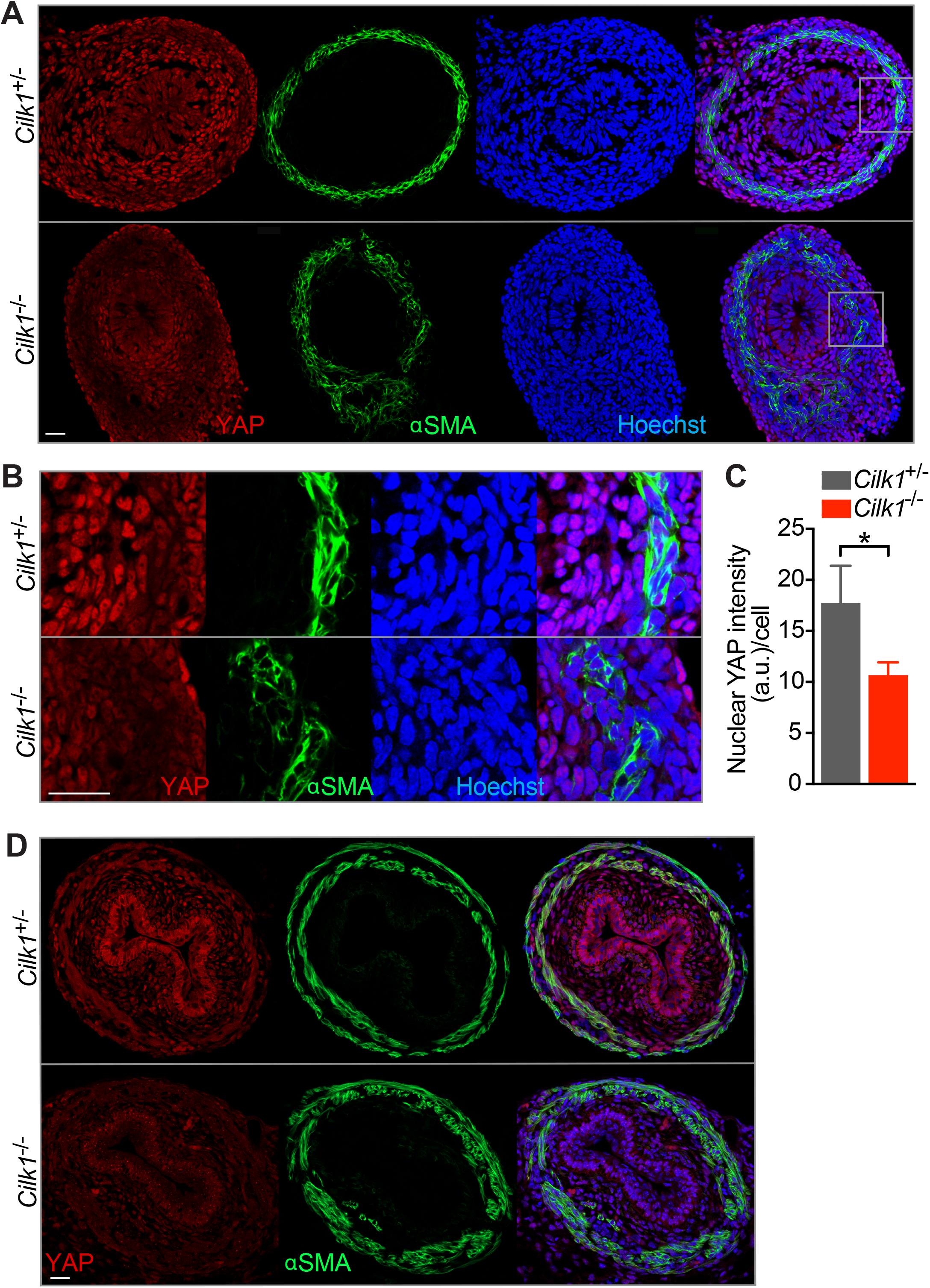
YAP levels depend on CILK1. (A) Immunofluorescence staining of E13.5 *Cilk1*^+/-^ and *Cilk1*^-/-^ intestines for YAP (red), smooth muscle (αSMA, green) and nuclei (Hoechst, blue). Scale bar, 25 μm. (B) Magnifications of the boxed areas from (A). Scale bar, 25 μm. (C) Nuclear YAP intensities of peri-epithelial mesenchymal cells in E13.5 *Cilk1*^+/-^ (n=4) and *Cilk1*^-/-^ (n=4) intestines. For each intestine, the nuclear YAP intensity per cell was averaged from three sections. Values are presented as means ± SEM. * p < 0.05 by ratio paired t test. (D) Immunofluorescence staining of E16.5 *Cilk1*^+/-^ and *Cilk1*^-/-^ esophagi for YAP (red), smooth muscle (αSMA, green) and nuclei (Hoechst, blue). Scale bar, 25 μm.

If YAP responds to the residual stress generated in the intestine by the circumferential smooth muscle, we predicted that disrupting the smooth muscle would decrease YAP nuclear levels. To test this prediction, we examined YAP localization in *Cilk1*^-/-^ intestine. Interestingly, the levels of nuclear YAP were decreased in *Cilk1*^-/-^ peri-epithelial stromal cells, indicating that CILK1 regulates YAP activity in the intestinal wall, directly or indirectly (Figure 7A-C). Similarly, in *Cilk1*^-/-^ esophagus, smooth muscle organization was disrupted, and nuclear YAP was reduced (Figure 7D).

Similarly, the levels of nuclear YAP were decreased in *Dermo*^Cre^ *Smo*^lox/lox^ peri-epithelial mesenchymal cells, revealing that mesenchymal HH signaling also promotes YAP nuclear levels (Figure S10A). Moreover, the levels of nuclear YAP were also decreased in *Myh11*^Cre-EGFP^ *R26*^DTA176/+^ peri-epithelial mesenchymal cells, demonstrating that intestinal smooth muscle is critical for YAP activation (Figure S10B). As mechanical strain activates YAP to drive cell proliferation in mammalian cells (Benham-Pyle et al., 2015), we hypothesized that YAP translates the residual stress generated by the smooth muscle into growth.

### Mesenchymal Hippo signaling promotes longitudinal intestinal growth

To test whether Hippo signaling participates in elongation, we generated *Dermo1*^Cre^ *Yap*^lox/lox^ *Taz*^lox/+^ embryos. *Dermo1*^Cre^ *Yap*^lox/lox^ *Taz*^lox/+^ embryos were grossly normal and exhibited no alteration in body size (Figure S11A). In addition, E13.5 *Dermo1*^Cre^ *Yap*^lox/lox^ *Taz*^lox/+^ intestines displayed no alteration in smooth muscle development (Figure S11B). Despite intact smooth muscle, *Dermo1*^Cre^ *Yap*^lox/lox^ *Taz*^lox/+^ intestines at E13.5 were 40% shorter than those of controls (Figure 8A-B), demonstrating that mesenchymal YAP activity is critical for elongation. *Dermo1*^Cre^ *Yap*^lox/+^ *Taz*^lox/lox^ intestines were 10% shorter than controls (Figure 8A-B), indicating that TAZ makes a more minor contribution to elongation. The finding, that YAP is dispensable for smooth muscle generation, but critical for elongation, is consistent with the hypothesis that smooth muscle activates YAP to promote growth.

**Figure 8.**
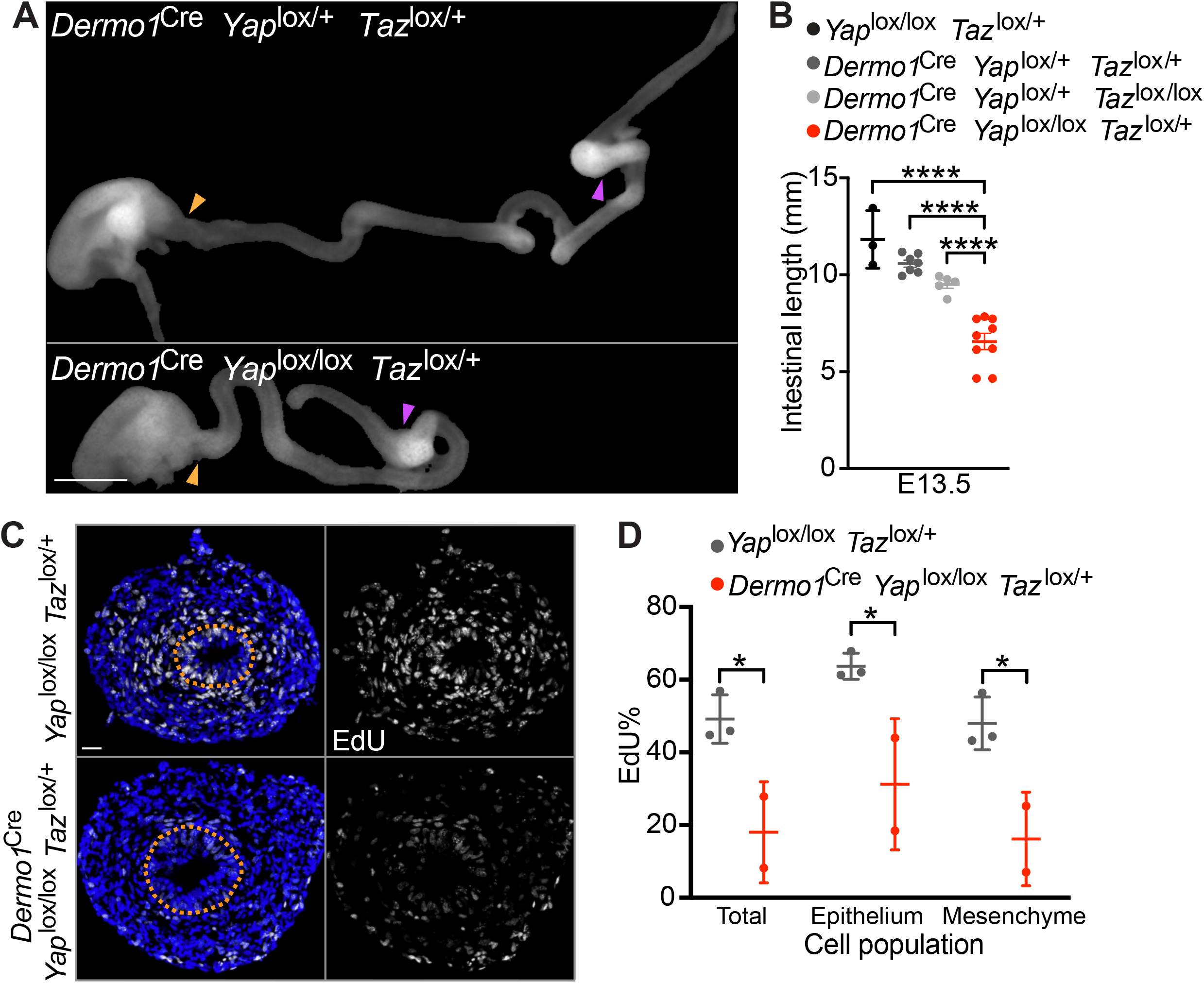
YAP functions in the mesenchyme to promote intestinal elongation and cell proliferation. (A) Photos of E13.5 *Dermo1*^Cre^ *YAP*^lox/+^ *TAZ*^lox/+^ and *Dermo1*^Cre^ *YAP*^lox/lox^ *TAZ*^lox/+^ intestines. Scale bar, 1 mm. The length of the small intestine was measured from the caudal stomach (orange arrowheads) to the rostral cecum (purple arrowheads). (B) Lengths of E13.5 in *YAP*^lox/lox^ *TAZ*^lox/+^, *Dermo1*^Cre^ *YAP*^lox/+^ *TAZ*^lox/lox^, *Dermo1*^Cre^ *YAP*^lox/+^ *TAZ*^lox/+^ and *Dermo1*^Cre^ *YAP*^lox/lox^ *TAZ*^lox/+^ intestines. Horizontal bars indicate means ± SD.**** p < 0.0001 by one-way ANOVA Sidak’s multiple comparisons test. (G)Immunofluorescence staining of E12.5 *YAP*^lox/lox^ *TAZ*^lox/+^ and *Dermo1*^Cre^ *YAP*^lox/lox^ *TAZ*^lox/+^ intestines for EdU (white) and nuclei (Hoechst, blue). Scale bar, 25 μm. Orange dotted lines outline the epithelium. (C) The percentage of EdU-positive cells in the E12.5 whole intestine, the epithelial cells or the mesenchymal cells in *YAP*^lox/lox^ *TAZ*^lox/+^ (n=3) and *Dermo1*^Cre^ *YAP*^lox/lox^ *TAZ*^lox/+^ (n=2). For each intestine, we averaged the percentages in the indicated cell population from three stained sections. Horizontal bars indicate means ± SD. *p < 0.05 by unpaired t test.

To examine whether mesenchymal YAP, like CILK1, promotes proliferation, we quantified EdU labeling of *Dermo1*^Cre^ *Yap*^lox/lox^ *Taz*^lox/+^ and control intestines at E12.5. The percentage of EdU-labeled cells was reduced in *Dermo1*^Cre^ *Yap*^lox/lox^ *Taz*^lox/+^ intestines both in the mesenchyme and epithelium (Figure 8C-D), indicating that the proliferation is is promoted by both CILK1 and YAP (Figure 2E-F). Thus, mesenchymal YAP is activated by mechanical forces generated by the circumferential smooth muscle and promotes proliferation to elongate the developing tubular organs.

## Discussion

Our results reveal a mechanism of tubular organ elongation by which intercellular signaling events create mechanical forces that are then translated into longitudinal growth. First, HH signals produced by epithelial cells are interpreted by the surrounding mesenchymal cells in a CILK1- and cilium-dependent manner. Second, ciliary HH signaling in the mesenchyme patterns the developing circumferential smooth muscle. Third, the smooth muscle produces residual stresses that exert mechanical forces on the mesenchymal cells. Fourth, these mesenchymal cells sense the smooth muscle-dependent forces, triggering nuclear accumulation of the Hippo pathway effector YAP. Fifth, nuclear YAP promotes cell proliferation, resulting in elongation. Thus, for the developing gut to elongate, morphogen-mediated patterning generates mechanical forces that drive YAP-promoted proliferation. As these findings apply to both the small intestine and the esophagus, we propose that the mechanical influences generated by ciliary signaling-patterned smooth muscle may be a general mechanism driving the elongation of tubular organs.

In support of the causal relationship between smooth muscle-generated forces and elongation, altering the organization or the number of circumferential smooth muscle cells by any of three independent means (disrupting ciliary signaling, disrupting HH signaling, or genetic ablation of smooth muscle cells) each reduces elongation. Importantly, disruption of smooth muscle decreases residual stress and the nuclear levels of YAP, and both presaged the emergence of decreased intestinal elongation. As mesenchymal YAP is essential for proliferation and elongation, we propose that the patterning of smooth muscle within the gut wall provides the mechanical growth cues which YAP interprets to drive elongation.

Many well studied cellular responses to mechanical forces are acute. For example, the cytoskeleton responds to forces within seconds (le Duc et al., 2010; Salbreux et al., 2012). Developmental processes typically occur over much longer timescales. In Drosophila, mechanical cues can affect epithelial planar organization to drive elongation (Stooke-Vaughan and Campàs, 2018; Vichas and Zallen, 2011). In the mammalian gut, our findings reveal that the smooth muscle-generated residual stress stores durable spatially-organized mechanical information. We propose that residual stress persists to enable tissue-level responses, such as gut elongation, over developmental timescales. Interestingly, mechanical cues can also be stored in extracellular matrix-based basement membrane as a stiffness gradient to drive elongation of the Drosophila follicle (Crest et al., 2017). Similar to our findings, mechanical forces activates YAP and elongates the Drosophila follicle, although proliferation is not involved (Fletcher et al., 2018). Static forces can activate YAP via deformation of the nucleus (Elosegui-Artola et al., 2017). Whether circumferential force activates YAP via nuclear stretching will be an interesting subject of further investigation.

The effects of mechanical forces in vertebrate tissue growth have been most extensively documented in bone remodeling in response to external forces (Christen et al., 2014; Vico et al., 2000). During development, our data suggest that mechanical forces, as in bone, promote growth, but these forces are internally generated by smooth muscle within the gut wall. Longitudinal forces transmitted by the vitelline duct also promote elongation of embryonic intestine in the chick embryo (Chevalier et al., 2018). The relative growth between the gut and its connecting tissues generates forces that shape the looping of gut tube across species (Savin et al., 2011), suggesting that organ elongation is regulated by both internal and external forces.

As mechanical force can promote elongation of tubular organs during development, we speculate that activation of mechanotransducive signaling may also promote regeneration in adult tissues. This speculation is supported by the emerging evidence showing that reversing the tissue mechanics associated with aging can rejuvenate the adult stem cells (Segel et al., 2019). The adult intestine is one of the most proliferative organs, with human intestinal epithelial cells turning over every 2-5 days (Darwich et al., 2014) driven by intestinal epithelial stem cells which demonstrate robust regenerative capacity also *in vitro* (Tuveson and Clevers, 2019; Wells and Spence, 2014). However, guts do not display compensatory lengthening postnatally, either after injury or congenital defects that cause shortening. Consequently, individuals with short bowel syndrome do not recover intestinal length over time, leading to chronic water and nutrient absorption deficits (Greig et al., 2019). Interestingly, application of longitudinal mechanical force can promote postnatal intestinal growth in animal models (Stark and Dunn, 2012). Our findings raise the possibility that application of radial forces may be therapeutically useful for promoting postnatal elongation of tubular organs.

We have found that at midgestation, the circumferential smooth muscle imparts tissue mechanical properties that induce proliferation and intestinal elongation. Later in the development, intestinal smooth muscle helps coordinate villus formation (Shyer et al., 2013). In addition, the spontaneous contractions of the same smooth muscle layer cues a second smooth muscle layer to align longitudinally along the intestine during late gestation (Huycke et al., 2019). Therefore, the mechanical force provided by smooth muscle participates at different times in multiple aspects of organ development.

Although both cilia and HH signaling are required for generating residual stress in the developing intestine, loss of ciliary proteins such as CILK alters the organization of the smooth muscle, whereas loss of SMO decreases the amount of smooth muscle. In both neural tube and limb bud development, loss of ciliary proteins and loss of SMO also lead to distinct phenotypes (Liu et al., 2005; Wijgerde et al., 2002). Activation of the HH signal transduction pathway is essential for conversion of GLI repressors into GLI activators, initiating the HH transcriptional program. Cilia are required for both GLI repressor and GLI activator formation. Thus, we propose that, at in other tissues, the different effects of loss of CILK and SMO on smooth muscle formation are attributable to attenuation of both GLI repressor and GLI activator in the former case, and attenuation only of GLI activator in the latter case.

We provided evidence that ciliary HH signaling in the intestinal mesenchyme patterns the circumferential smooth muscle to promote growth. The role of HH signaling in intestinal growth is therefore distinct from its roles in tissues such as the cerebellum and skin, where it directly induces the expression of drivers of the cell cycle (Lopez-Rios et al., 2012; Raleigh et al., 2018). Instead, the intestine represents a distinct model of how HH signaling functions in organ development: HH signaling patterns the gut, and this patterning, acting via mechanical influences interpreted by the Hippo pathway, promotes proliferation.

To conclude, we uncovered an integrated molecular mechanism by which morphogen signaling patterns a tissue to create the mechanical cues that regulate organ size. During the development of the tubular organs, the activation of ciliary signaling in the mesenchyme patterns the wrapping smooth muscle layer, which generates intrinsic forces that activates the mechanotransducive effector YAP. Active YAP promotes proliferation resulting in growth. This cooperation between biochemical and mechanical signals to regulate organ elongation may illustrated here represent a general mechanism to control growth of tubular organs.

## Acknowledgements

We are grateful to Elaine Carlson for help in generating *Tctn3* mutant alleles and E Yu for husbandry and genotyping of the mice used in this study. We thank Amy E. Shyer, Tyler R. Huycke, Wallace Marshall, Zev J. Gartner and Ophir D. Klein for thoughtful discussions. We also thank the members of the Reiter lab and the Makela lab for critical comments and suggestions on the study. This work was funded by NIH NIGMS R01GM095941, R01AR054396, and NICHD R01HD089918 to J.F.R., NIH P30 DK090868 to K.E.M., and UCSF Program for Breakthrough Biomedical Research (PBBR) to K.E.M. and J.F.R..

## Author contributions

J.F.R. and Y.Y. conceived the study and designed the experiments, except for the observations in Figure 1A, 1B, 1C, S1B, S1C, S1D, S3 and S4, which were initially made by Y.Y. and P.P. with T.P.M. Y.Y. with K.E.M. and J.F.R made the findings in Figure 1D, 1E, 1F, 2A, S2B, S2C, and S5. P.P. generated the data in Figure S4 and repeated the results in Figure 1A, 1B, 2A, S1B, S5B independently. C.X. generated the data in Figure 2B. A.L.K. generated the *Tctn3* mutant mouse line. Y.Y. performed all the other experiments. J.F.R and Y.Y. wrote and prepared the manuscript with input from all authors.

## Figure Legends

**Figure S1.**
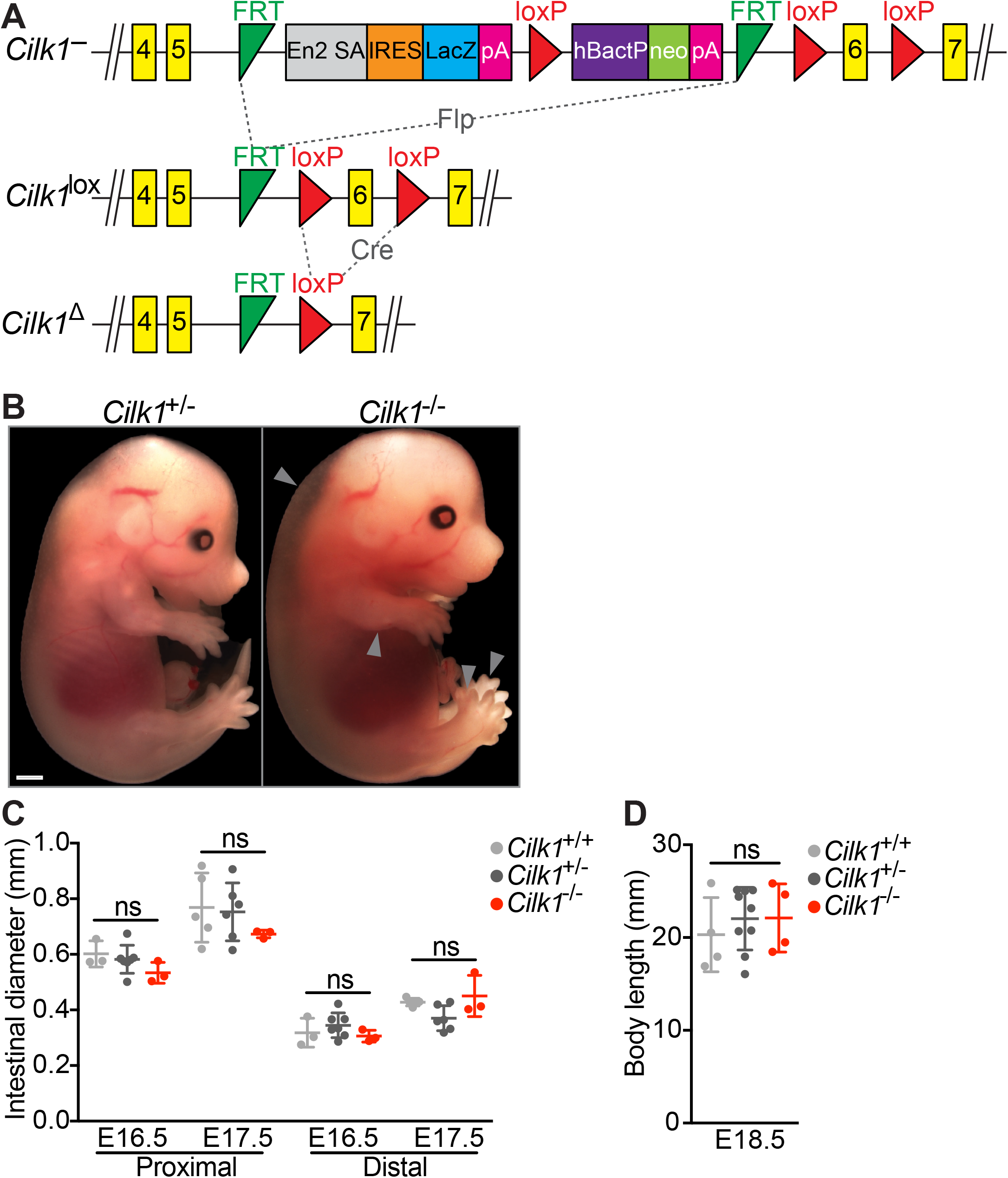
CILK1 is essential for limb patterning, but does not affect intestinal diameter or the whole-body length. Related to Figure 1. (A) Schematic representation of the murine *Cilk1* alleles in this study. The *Cilk1*^-^ allele, also called *Cilk1*^tm1a(KOMP)Mbp^, is a “knockout first” allele that contains a cassette expressing *lacZ* gene and *neomycin resistance gene* (neo) inserted between exon 5 and exon 6 of *Cilk1*. Flp-mediated recombination removed the cassette between the FRT sites to generate the conditional allele, *Cilk1*^lox^. *Cilk1*^lox^ possesses exon 6 flanked by loxP sites. Cre-mediated recombination removed exon 6 to generate a deletion allele, *Cilk1*^Δ^. FRT, Flippase recognition target; En2 SA, splice acceptor of mouse *Engrailed2* exon 2; IRES, an internal ribosome entry sequence, pA, polyadenylation signal (pA); hBactP, human *beta-actin* promoter. Only exons 4-7 of the *Cilk1* gene are shown. (B) Photos of E15.5 *Cilk1*^+/-^ and *Cilk1*^-/-^ embryos. Arrowheads indicate edema, short limbs and polydactyly. Scale bar, 1 mm. (C) Diameters of E16.5 and E17.5 *Cilk1^+/+^, Cilk1*^+/-^ and *Cilk1*^-/-^ proximal and distal intestines. (D) The crown-rump lengths of E18.5 *Cilk1^+/+^, Cilk1*^+/-^ and *Cilk1*^-/-^ embryos. (C-D) Each point in the scatter plots represents the value from an individual embryo. Horizontal bars indicate means ± SD. ns, p > 0.05 by ordinary one-way ANOVA test performed for each embryonic stage.

**Figure S2.**
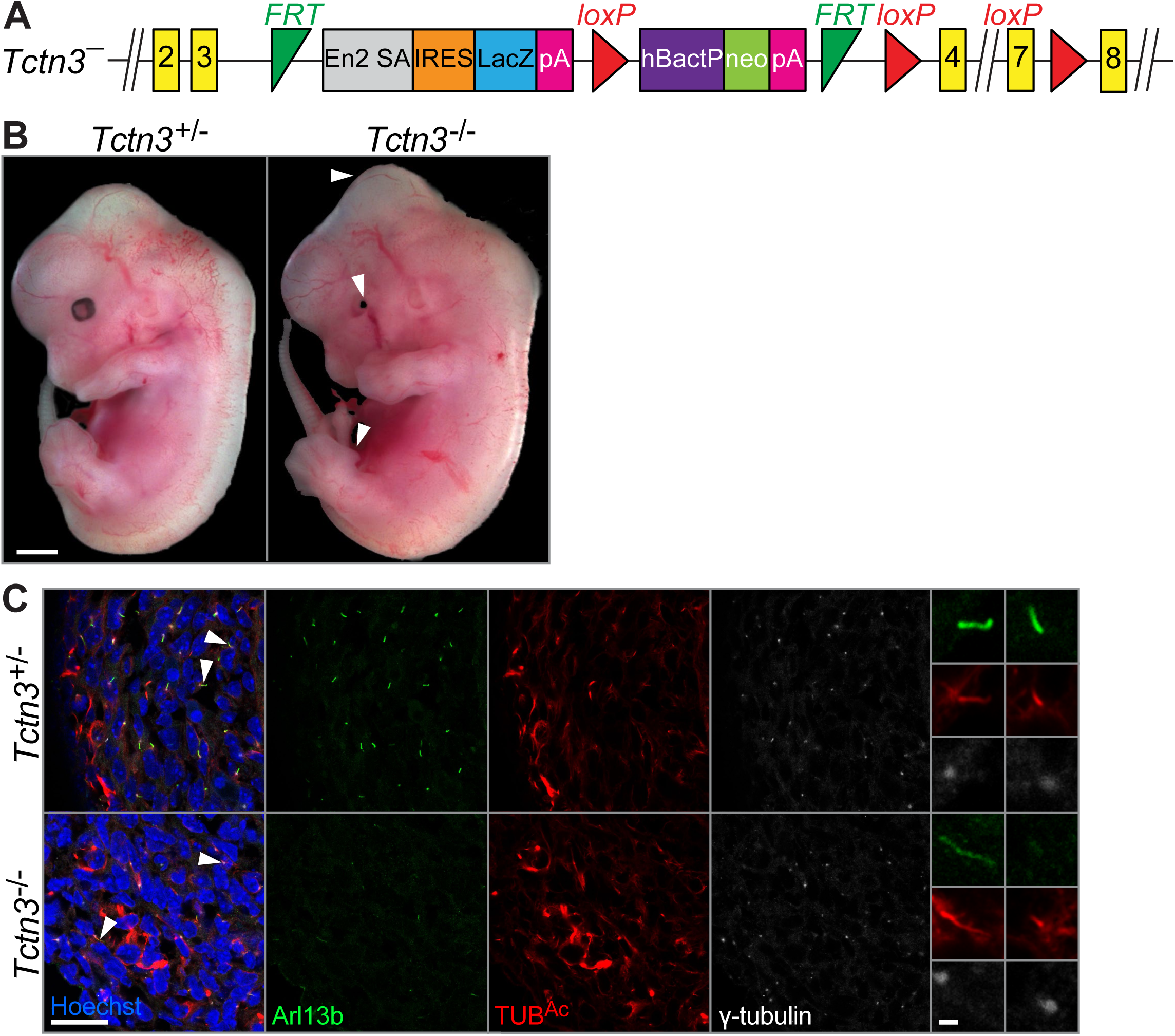
TCTN3 is required for developmental patterning and ciliary localization of ARL13B. Related to Figure 1. (A) Schematic representation of the murine *Tctn3* alleles in this study. The *Tctn3*^-^ allele, also called *Tctn3*^tm47188(L1L2_Bact_P)^, contains a cassette expressing *lacZ* gene and *neomycin resistance gene* (neo) inserted between exon 3 and exon 4. FRT, flippase recognition target; En2 SA, splice acceptor of mouse *Engrailed2* exon 2; IRES, an internal ribosome entry sequence, pA, polyadenylation (pA); hBactP, human beta actin promoter. Only exons 2-8 of the *Tctn3* gene are shown. (B) Photos of E13.5 embryos *Tctn3*^+/-^ and *Tctn3*^-/-^. Arrowheads indicate hydrocephalus, microphthalmia and polydactyly. Scale bar, 1 mm. (C) Immunofluorescence staining for ciliary membrane (ARL13B, green), ciliary axonemes (TUB^Ac^, red), basal bodies (γ-tubulin, white) and nuclei (Hoechst, blue), in intestines of *Tctn3*^+/-^ and *Tctn3*^-/-^ embryos at E13.5. Scale bar, 25 μm. Scale bar for magnifications of the indicated cilia (right), 1 μm.

**Figure S3.**
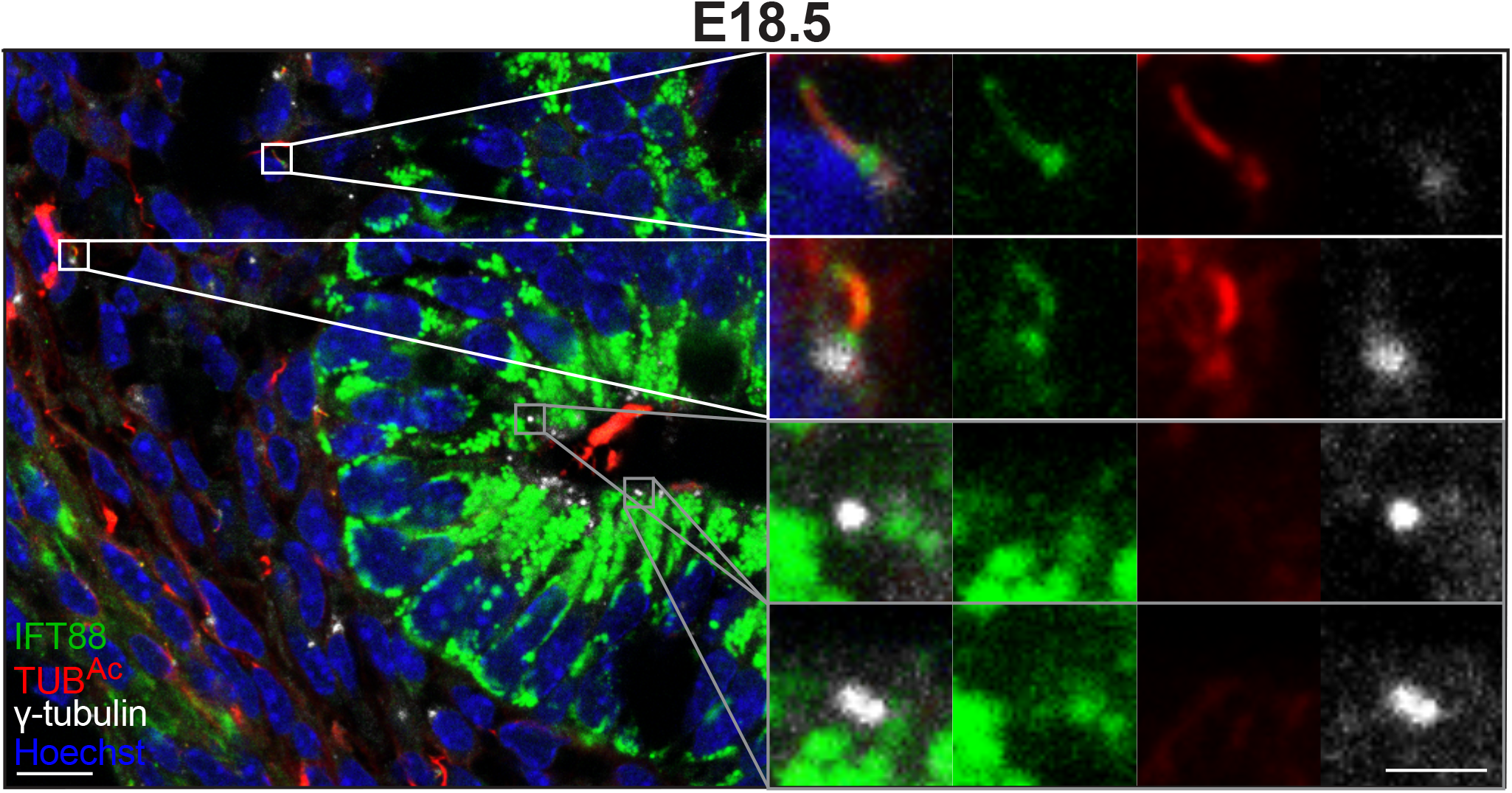
Intestinal mesenchymal cells, but not epithelial cells, possess cilia at E18.5. Related to Figure 2. Immunofluorescence staining for IFT88 (green), ciliary axonemes (TUB^Ac^, red), basal bodies (γ-tubulin, white) and nuclei (Hoechst, blue) in E18.5 intestines. Scale bar, 10 μm. White boxes denote mesenchymal cilia. Grey boxes denote epithelial centrosomes. Scale bar for magnifications, 2 μm.

**Figure S4.**
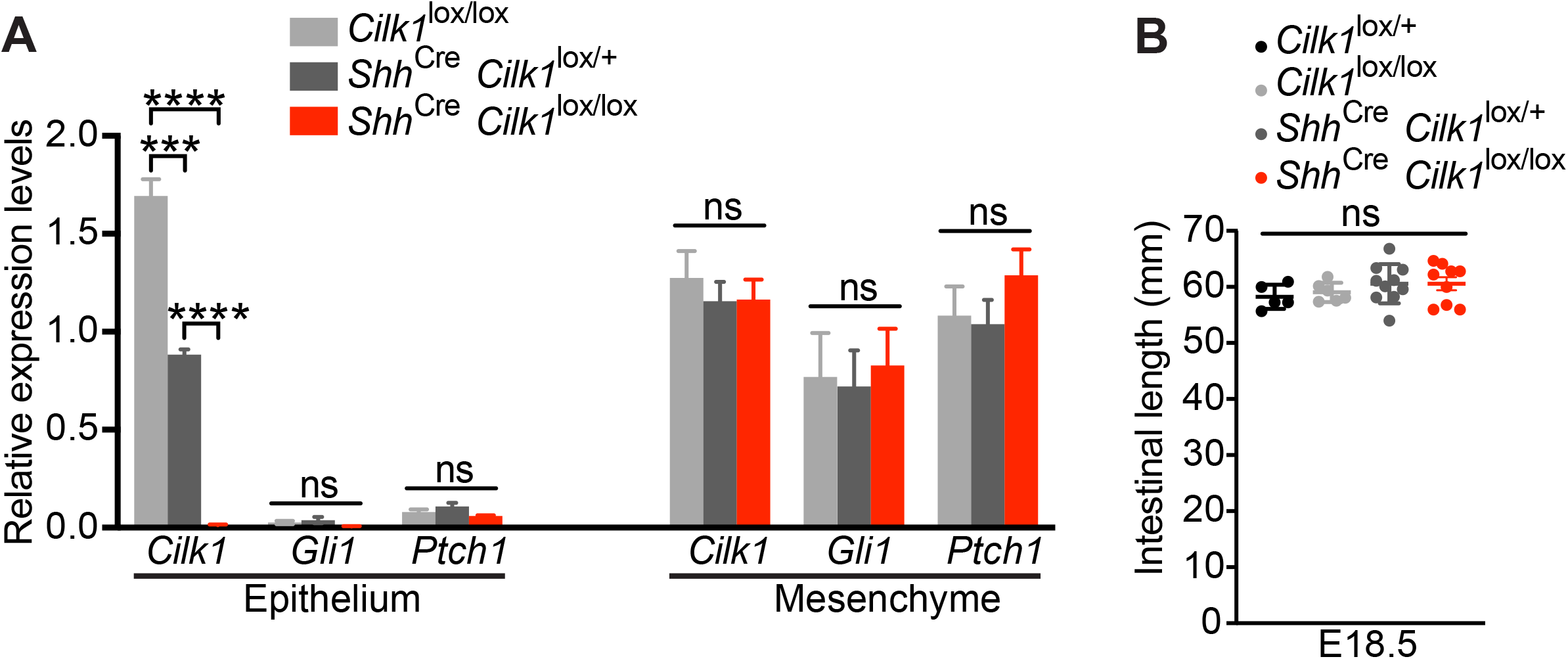
Epithelial CILK1 is dispensable for expression of HH target genes and intestinal elongation. Related to Figure 2. (A) Relative levels of *Cilk1, Gli1* and *Ptch1* mRNA in E18.5 intestines of control (*Cilk1*^lox/lox^, n=5 and *Shh*^Cre^ *Cilk1*^lox/+^, n=8) and *Shh*^Cre^ *Cilk1*^lox/lox^ (n=5) embryos. Values are presented as means ± SEM. ns, p > 0.05; *** p < 0.001; **** p < 0.0001 by two-way ANOVA Tukey’s multiple comparisons test. (B) Lengths of the small intestines in E18.5 control (*Cilk1*^lox/+^, n=5, *Cilk1*^lox/lox^, n=6 and *Shh*^Cre^ *Cilk1*^lox/+^, n=10) and *Shh*^Cre^ *Cilk1*^lox/lox^ (n=9) embryos. Each point in the scatter plots represents the value from an individual embryo. Horizontal bars indicate means ± SD. ns, p > 0.05 by ordinary one-way ANOVA test.

**Figure S5.**
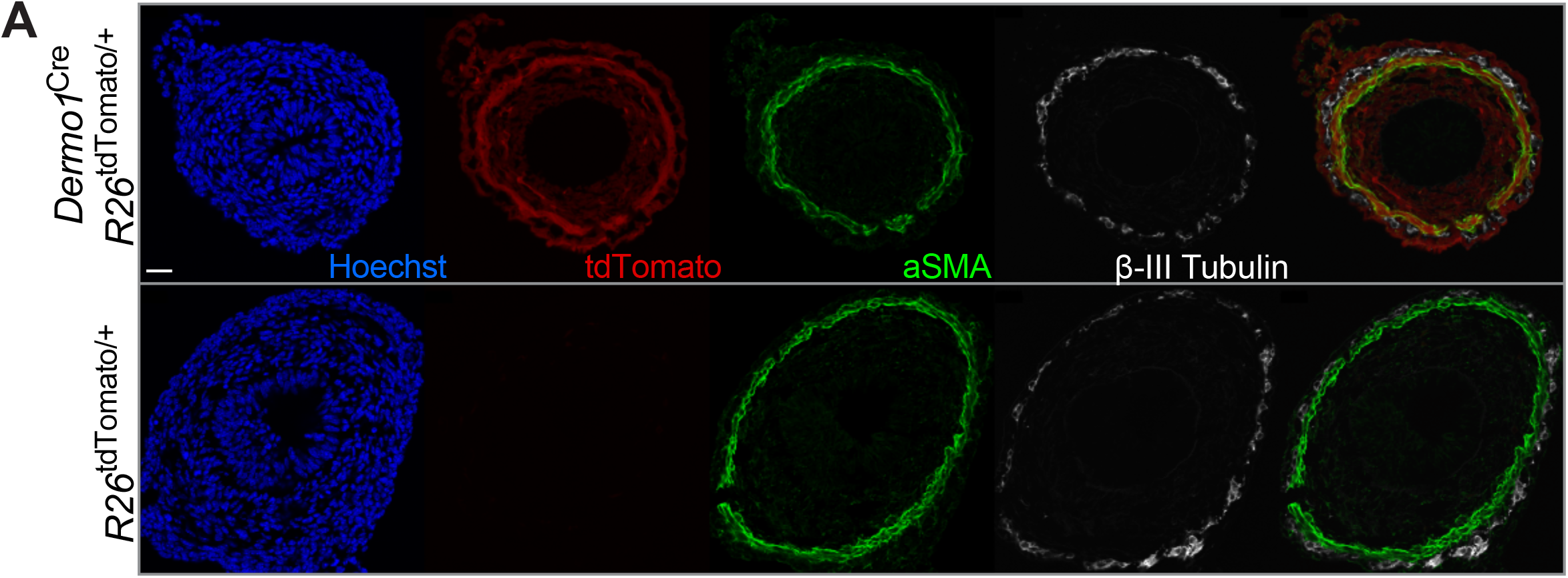
*Dermo1*^Cre^ mediates efficient recombination in the intestinal mesenchyme. Related to Figure 2. (A) Immunofluorescence staining of *Dermo1*^Cre^ *R26*^tdTomato/+^ and *R26*^tdTomato/+^ intestine at E13.5 for recombined cells (tdTomato, red), smooth muscle (αSMA, green), enteric nervous system (β-III Tubulin, white) and nuclei (Hoechst, blue). Scale bar, 25 μm. The endoderm-derived epithelial cells and the ectoderm-derived enteric nervous system cells do not express tdTomato. In contrast, mesoderm-derived mesenchymal cells, including the αSMA cells, express tdTomato.

**Figure S6.**
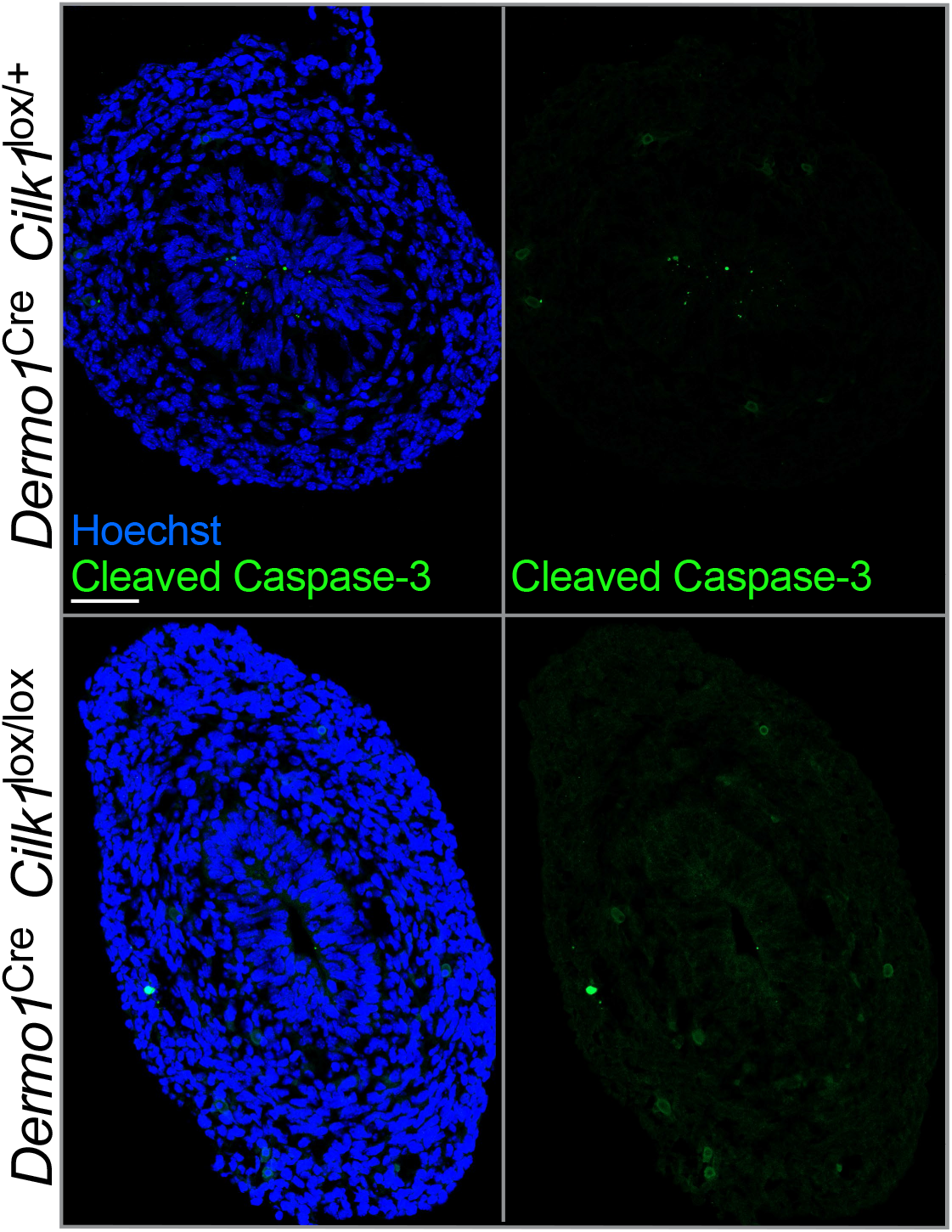
No increased apoptotic cells in *Dermo1*^Cre^ *Cilk1*^lox/lox^. Related to Figure 2. Immunofluorescence staining of Cleaved Caspase-3 (green) in E13.5 control *Dermo1*^Cre^ *Cilk1*^lox/+^ and *Dermo1*^Cre^ *Cilk1*^lox/lox^ intestines. Scale bar, 50 μm.

**Figure S7.**
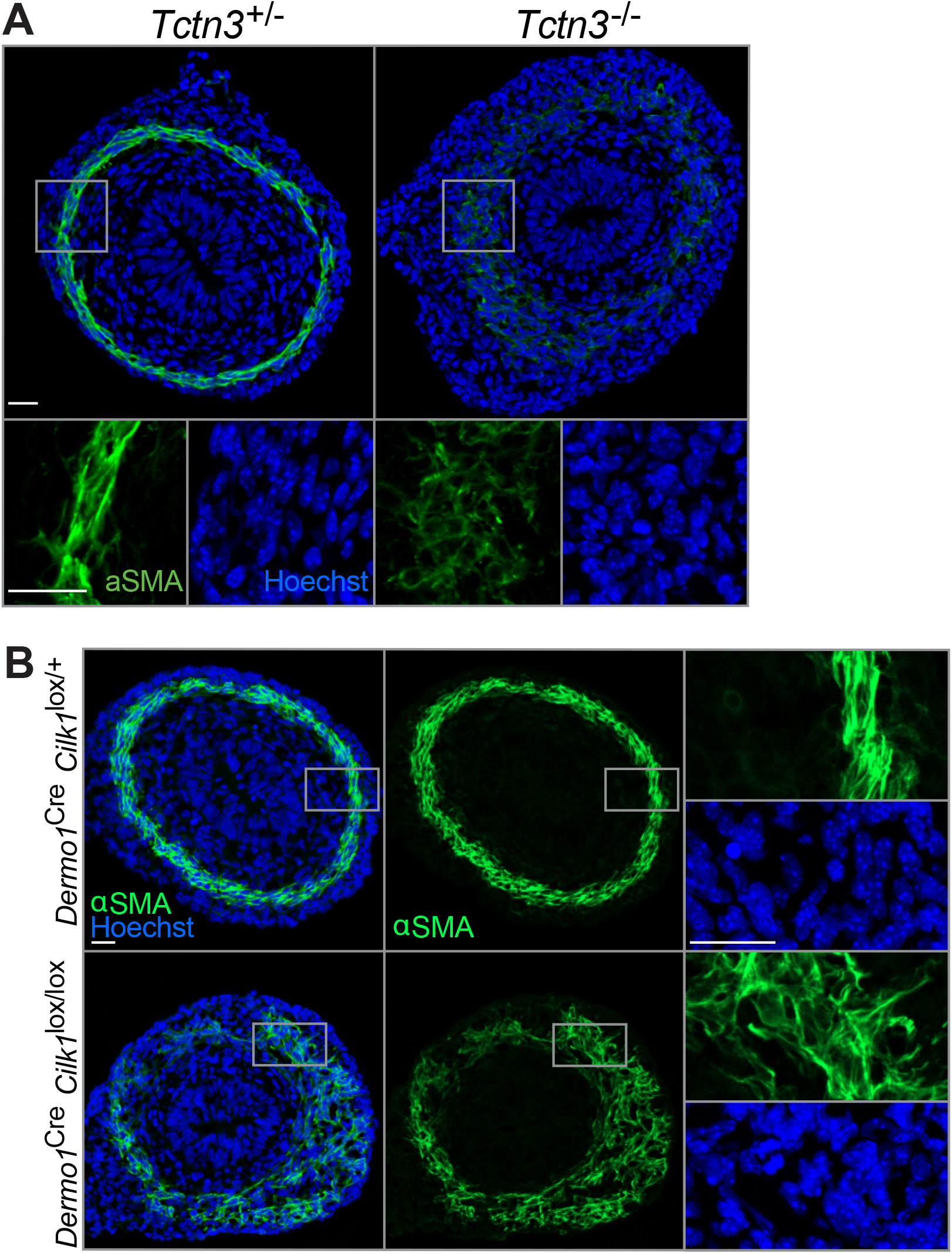
TCTN3 and mesenchymal CILK1 are required for circumferential smooth muscle organization. Related to Figure 3. (A) Immunofluorescence staining for smooth muscle (αSMA, green) and nuclei (Hoechst, blue) in E13.5 *Tctn3*^+/-^ and *Tctn3*^-/-^ intestines. Below, higher magnifications of boxed regions. Scale bars, 25 μm. (B) mmunofluorescence staining for smooth muscle (αSMA, green) and nuclei (Hoechst, blue) in E13.5 *Dermo1*^Cre^ *Cilk1*^lox/+^ and *Dermo1*^Cre^ *Cilk1*^lox/lox^ intestines. Right, higher magnifications of boxed regions. Scale bars, 25 μm.

**Figure S8.**
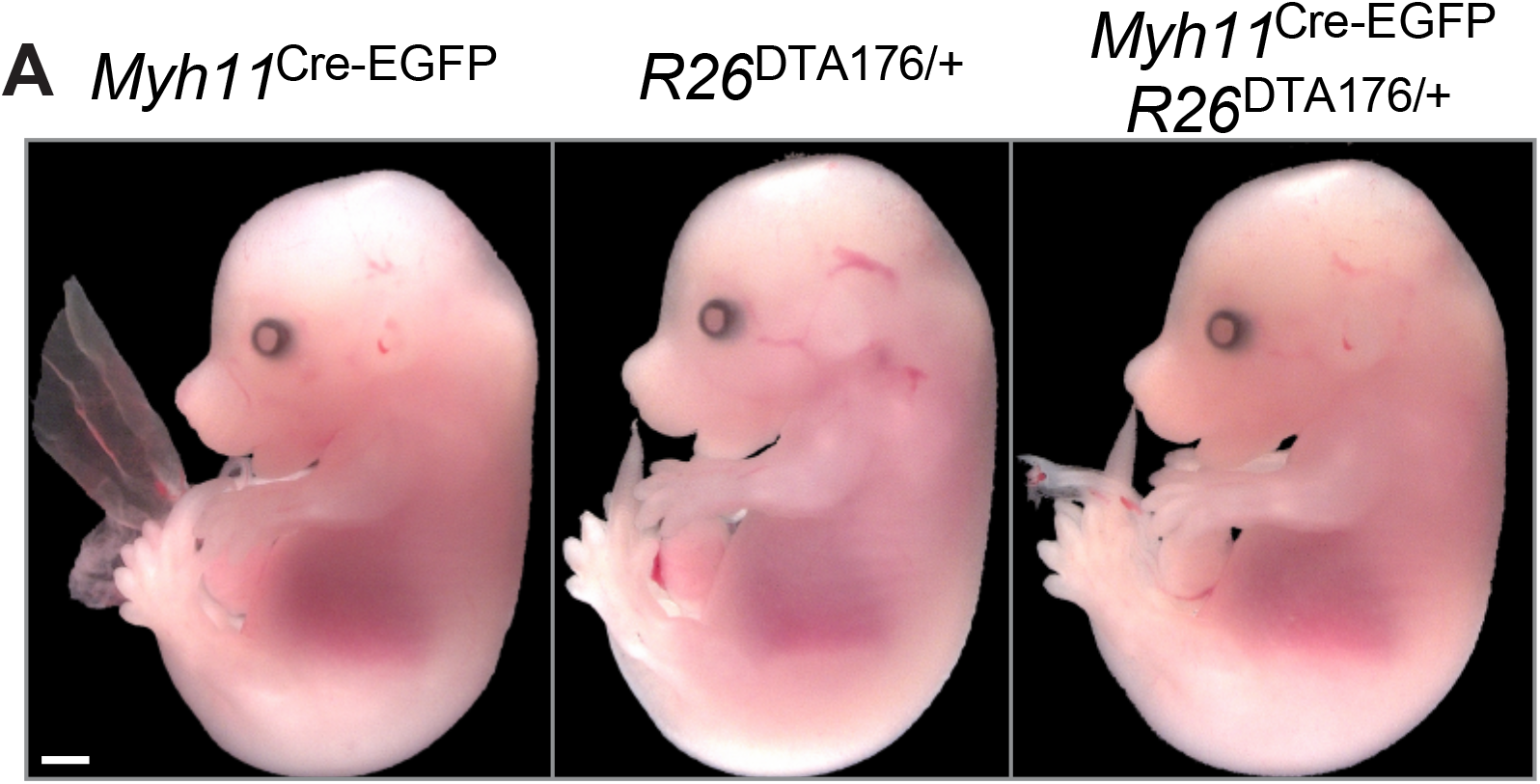
Partially ablating smooth muscle cells does not grossly disrupt embryogenesis. Related to Figure 5. Photos of E15.5 control (*Myh11*^Cre-EGFP^ and *R26*^DTA176/+^) and *Myh11*^Cre-EGFP^ *R26*^DTA176/+^ embryos. Scale bar, 1 mm.

**Figure S9.**
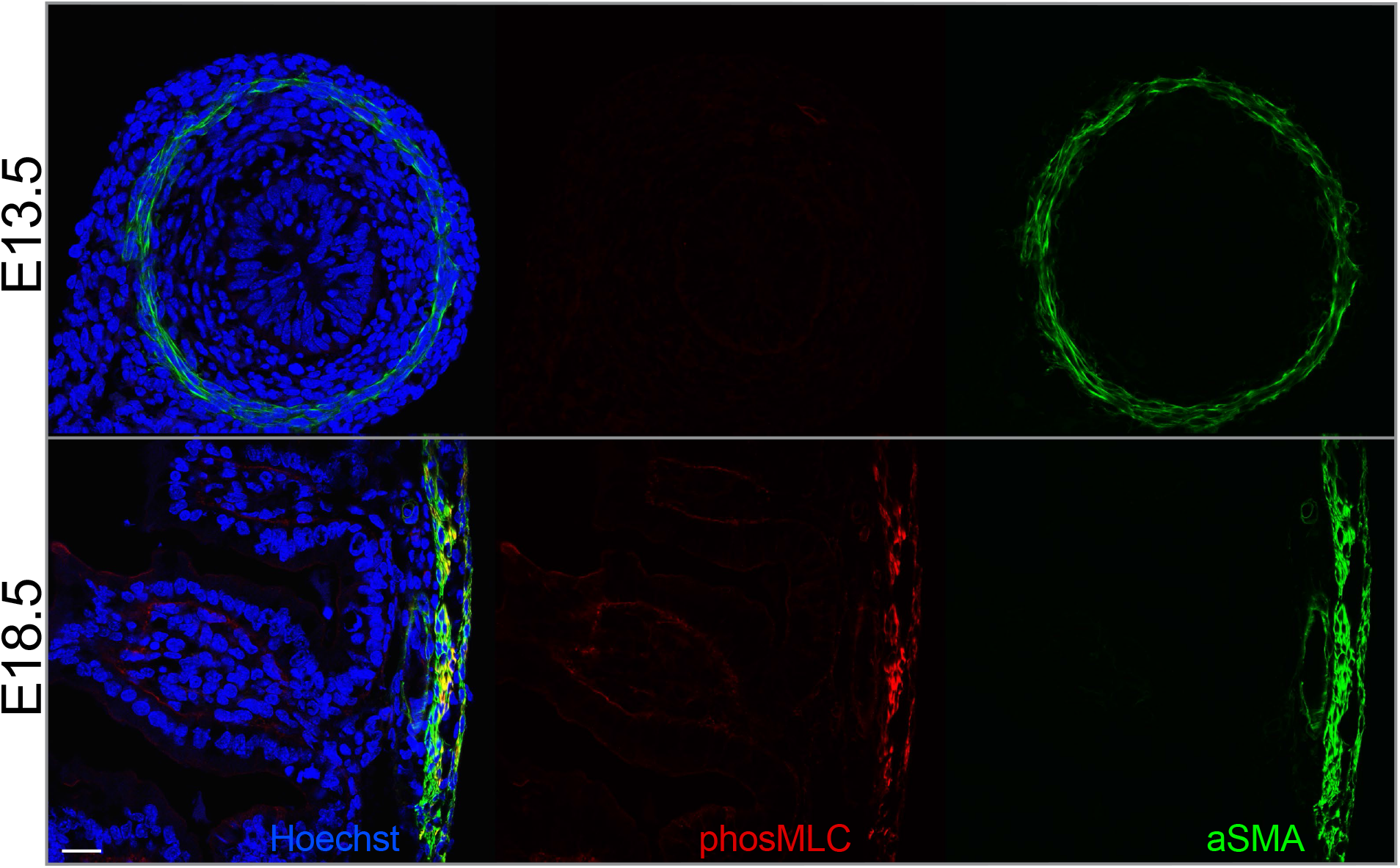
The smooth muscle cells are negative for phos-MLC at E13.5. Related to Figure 6. Immunofluorescence staining of E13.5 and E18.5 wild-type intestines for phospho-MLC (red), smooth muscle (αSMA, green) and nuclei (Hoechst, blue). Scale bar, 25 μm.

**Figure S10.**
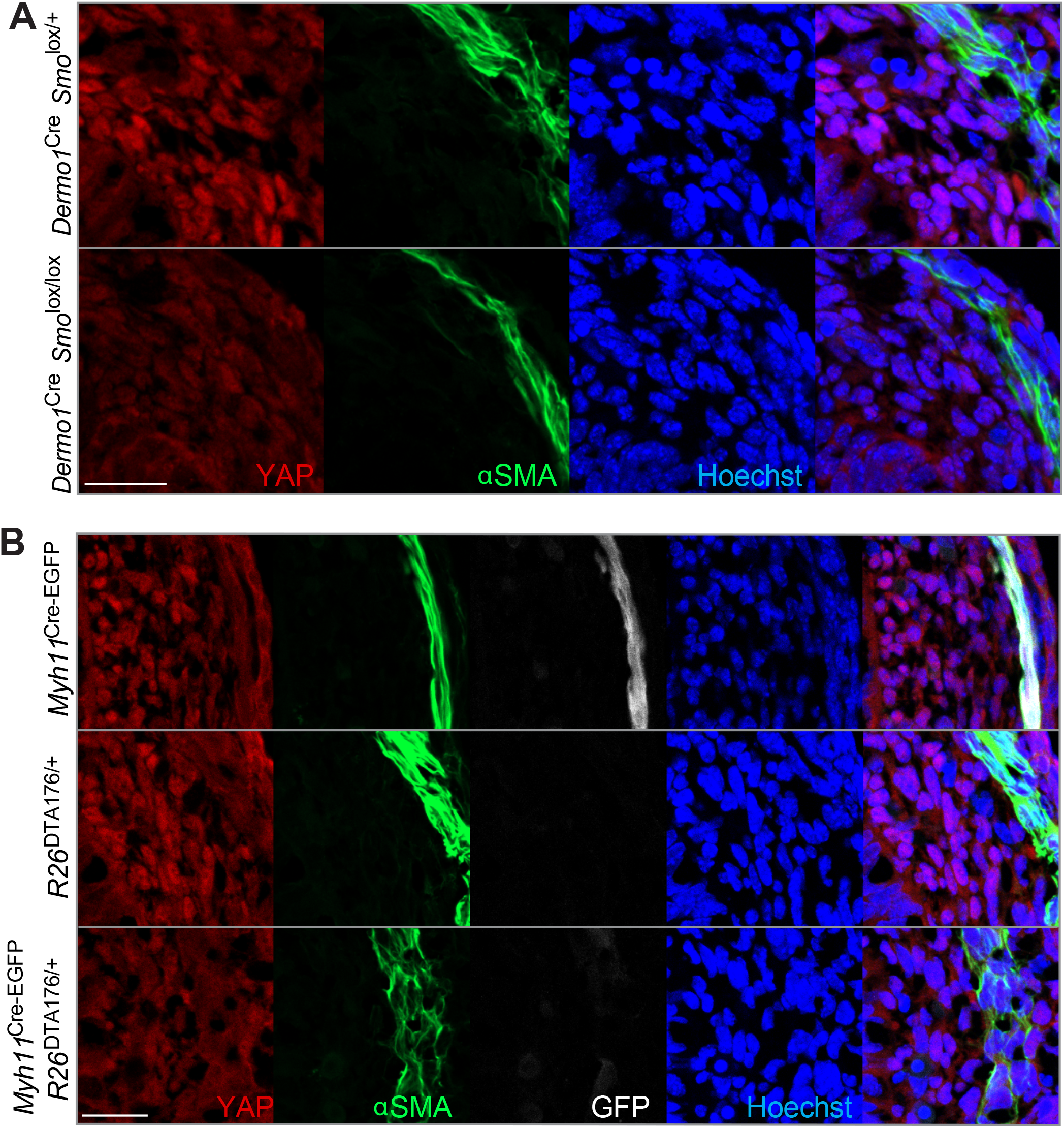
Nuclear levels of YAP depend on SMO function in the mesenchyme and smooth muscle integrity. Related to Figure 7. (A) Immunofluorescence staining for YAP (red), smooth muscle (αSMA, green) and nuclei (Hoechst, blue) in E13.5 *Dermo1*^Cre^ *Smo*^lox/+^ and *Dermo1*^Cre^ *Smo*^lox/lox^ intestines. Scale bar, 25 μm. (B) Immunofluorescence staining for YAP (red), smooth muscle (αSMA, green) and nuclei (Hoechst, blue) in E14.5 control (*Myh11*^Cre-EGFP^ and *R26*^DTA176/+^) and *Myh11*^Cre-EGFP^ *R26*^DTA176/+^ intestines. Scale bar, 25 μm.

**Figure S11.**
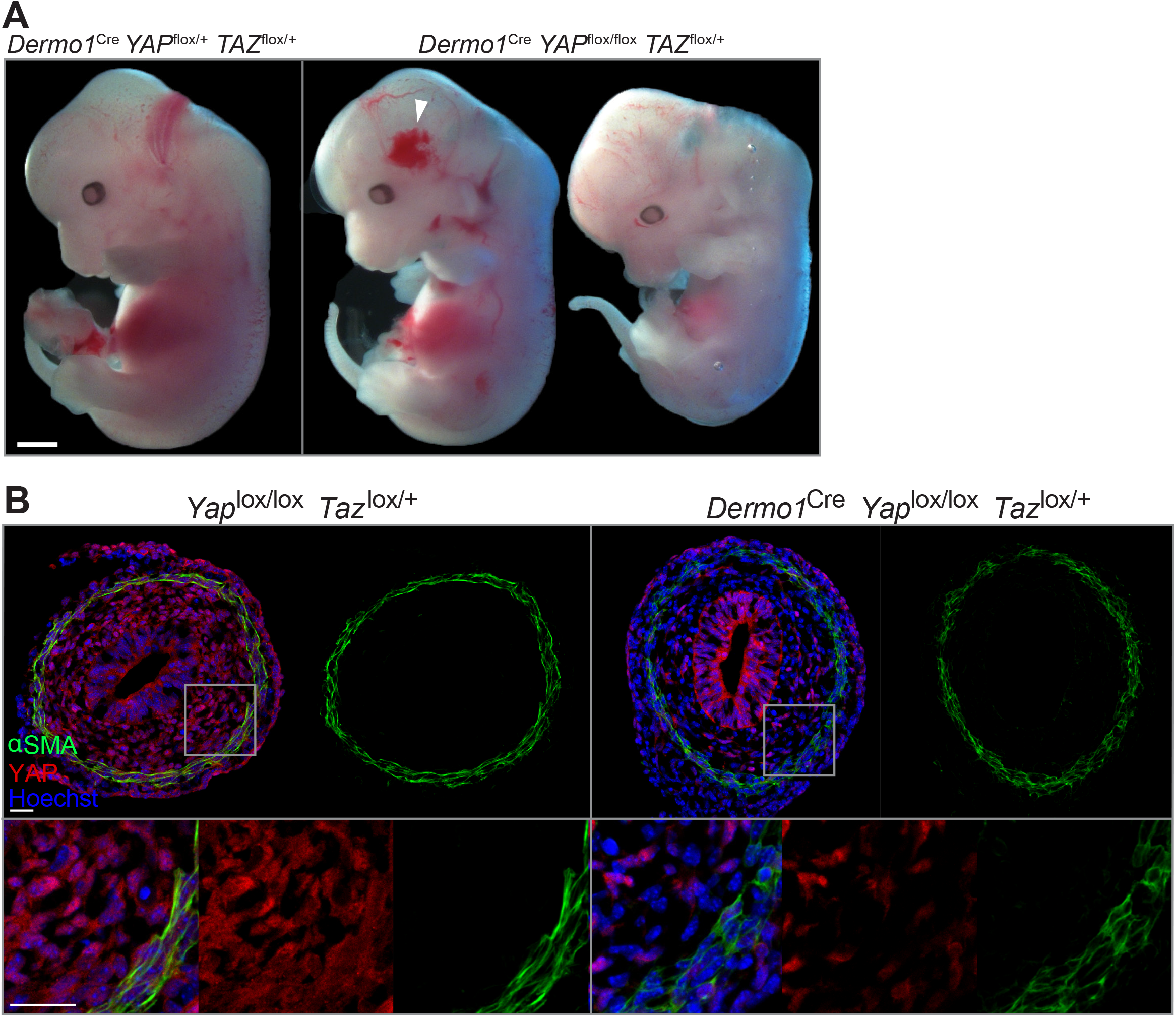
Deletion of YAP in the mesenchyme causes semi-penetrant hemorrhage and does not affect whole-body size or development of the circumferential smooth muscle at E13.5. (A) Photos of E13.5 *Dermo1*^Cre^ *YAP*^lox/+^ *TAZ*^lox/+^ and *Dermo1*^Cre^ *YAP*^lox/lox^ *TAZ*^lox/+^ embryos. Arrowhead indicates hemorrhage in one of the *Dermo1*^Cre^ *YAP*^lox/lox^ *TAZ*^lox/+^ embryos. Scale bar, 1 mm. (B) Immunofluorescence staining of E13.5 *YAP^lox/+^ TAZ*^lox/lox^ and *Dermo1*^Cre^ *YAP^lox/lox^ TAZ*^lox/+^ intestines for YAP (red), smooth muscle (αSMA, green) and nuclei (Hoechst, blue). Below, higher magnifications of boxed regions. Scale bars, 25 μm.

## References

Allen, B.G., and Walsh, M.P. (1994). The biochemical basis of the regulation of smoothmuscle contraction. Trends in Biochemical Sciences 19, 362–368.

Bangs, F., and Anderson, K.V. (2017). Primary Cilia and Mammalian Hedgehog Signaling. Cold Spring Harb Perspect Biol 9, a028175.

Benham-Pyle, B.W., Pruitt, B.L., and Nelson, W.J. (2015). Cell adhesion. Mechanical strain induces E-cadherin-dependent Yap1 and β-catenin activation to drive cell cycle entry. Science 348, 1024–1027.

Chaya, T., Omori, Y., Kuwahara, R., and Furukawa, T. (2014). ICK is essential for cell type-specific ciliogenesis and the regulation of ciliary transport. Embo J. 33, 1227–1242.

Chávez, M., Ena, S., Van Sande, J., de Kerchove d’Exaerde, A., Schurmans, S., and Schiffmann, S.N. (2015). Modulation of Ciliary Phosphoinositide Content Regulates Trafficking and Sonic Hedgehog Signaling Output. Developmental Cell 34, 338–350.

Chevalier, N.R., de Witte, T.-M., Cornelissen, A.J.M., Dufour, S., Proux-Gillardeaux, V., and Asnacios, A. (2018). Mechanical Tension Drives Elongational Growth of the Embryonic Gut. Sci Rep 8, 5995.

Chin, A.M., Hill, D.R., Aurora, M., and Spence, J.R. (2017). Morphogenesis and maturation of the embryonic and postnatal intestine. Semin. Cell Dev. Biol. 66, 81–93.

Christen, P., Ito, K., Ellouz, R., Boutroy, S., Sornay-Rendu, E., Chapurlat, R.D., and van Rietbergen, B. (2014). Bone remodelling in humans is load-driven but not lazy. Nature Communications 5, 4855–4855.

Corbit, K.C., Aanstad, P., Singla, V., Norman, A.R., Stainier, D.Y.R., and Reiter, J.F. (2005). Vertebrate Smoothened functions at the primary cilium. Nature 437, 1018–1021.

Cotton, J.L., Li, Q., Ma, L., Park, J.-S., Wang, J., Ou, J., Zhu, L.J., Ip, Y.T., Johnson, R.L., and Mao, J. (2017). YAP/TAZ and Hedgehog Coordinate Growth and Patterning in Gastrointestinal Mesenchyme. Developmental Cell 43, 35–47.e4.

Crest, J., Diz-Munoz, A., Chen, D.Y., Fletcher, D.A., and Bilder, D. (2017). Organ sculpting by patterned extracellular matrix stiffness. Elife 6, 773.

Darwich, A.S., Aslam, U., Ashcroft, D.M., and Rostami-Hodjegan, A. (2014). Meta-Analysis of the Turnover of Intestinal Epithelia in Preclinical Animal Species and Humans. Drug Metab Dispos 42, 2016–2022.

Dong, J., Feldmann, G., Huang, J., Wu, S., Zhang, N., Comerford, S.A., Gayyed, M.F., Anders, R.A., Maitra, A., and Pan, D. (2007). Elucidation of a universal size-control mechanism in Drosophila and mammals. Cell 130, 1120–1133.

Dowdle, W.E., Robinson, J.F., Kneist, A., Sirerol-Piquer, M.S., Frints, S.G.M., Corbit, K.C., Zaghloul, N.A., Zaghloul, N.A., van Lijnschoten, G., Mulders, L., et al. (2011). Disruption of a ciliary B9 protein complex causes Meckel syndrome. Am. J. Hum. Genet. 89, 94–110.

Dupont, S., Morsut, L., Aragona, M., Enzo, E., Giulitti, S., Cordenonsi, M., Zanconato, F., Le Digabel, J., Forcato, M., Bicciato, S., et al. (2011). Role of YAP/TAZ in mechanotransduction. Nature 474, 179–183.

Dyson, J.M., Conduit, S.E., Feeney, S.J., Hakim, S., DiTommaso, T., Fulcher, A.J., Sriratana, A., Ramm, G., Horan, K.A., Gurung, R., et al. (2017). INPP5E regulates phosphoinositide-dependent cilia transition zone function. J. Cell Biol. 216, 247–263.

Elosegui-Artola, A., Andreu, I., Beedle, A.E.M., Lezamiz, A., Uroz, M., Kosmalska, A.J., Oria, R., Kechagia, J.Z., Rico-Lastres, P., Le Roux, A.-L., et al. (2017). Force Triggers YAP Nuclear Entry by Regulating Transport across Nuclear Pores. Cell 171, 1397–1410.e14.

FitzSimmons, J., Chinn, A., and Shepard, T.H. (1988). Normal length of the human fetal gastrointestinal tract. Pediatr Pathol 8, 633–641.

Fletcher, G.C., Diaz-de-la-Loza, M.-D.-C., Borreguero-Muñoz, N., Holder, M., Aguilar-Aragon, M., and Thompson, B.J. (2018). Mechanical strain regulates the Hippo pathway in Drosophila. Development 145, dev159467.

Fu, Z., Gailey, C.D., Wang, E.J., and Brautigan, D.L. (2019). Ciliogenesis associated kinase 1: targets and functions in various organ systems. FEBS Lett. 593, 2990–3002.

Fung, Y.C. (1991). What are the residual stresses doing in our blood vessels? Ann Biomed Eng 19, 237–249.

Garcia-Gonzalo, F.R., Corbit, K.C., Sirerol-Piquer, M.S., Ramaswami, G., Otto, E.A., Noriega, T.R., Seol, A.D., Robinson, J.F., Bennett, C.L., Josifova, D.J., et al. (2011). A transition zone complex regulates mammalian ciliogenesis and ciliary membrane composition. Nat. Genet. 43, 776–784.

Garcia-Gonzalo, F.R., Phua, S.C., Roberson, E.C., Garcia, G., Abedin, M., Schurmans, S., Inoue, T., and Reiter, J.F. (2015). Phosphoinositides Regulate Ciliary Protein Trafficking to Modulate Hedgehog Signaling. Developmental Cell 34, 400–409.

García-García, M.J., Eggenschwiler, J.T., Caspary, T., Alcorn, H.L., Wyler, M.R., Huangfu, D., Rakeman, A.S., Lee, J.D., Feinberg, E.H., Timmer, J.R., et al. (2005). Analysis of mouse embryonic patterning and morphogenesis by forward genetics. Proceedings of the National Academy of Sciences 102, 5913–5919.

Gregersen, H., and Kassab, G. (1996). Biomechanics of the gastrointestinal tract. Neurogastroenterol Motil 8, 277–297.

Greig, C.J., Oh, P.S., Gross, E.R., and Cowles, R.A. (2019). Retracing our STEPs: Four decades of progress in intestinal lengthening procedures for short bowel syndrome. The American Journal of Surgery 217, 772–782.

Hakem, R., Hakem, A., Duncan, G.S., Henderson, J.T., Woo, M., Soengas, M.S., Elia, A., la Pompa, de, J.L., Kagi, D., Khoo, W., et al. (1998). Differential requirement for caspase 9 in apoptotic pathways in vivo. Cell 94, 339–352.

Han, H.C., and Fung, Y.C. (1991). Residual strains in porcine and canine trachea. Journal of Biomechanics 24, 307–315.

Harfe, B.D., Scherz, P.J., Nissim, S., Tian, H., McMahon, A.P., and Tabin, C.J. (2004). Evidence for an expansion-based temporal Shh gradient in specifying vertebrate digit identities. Cell 118, 517–528.

Haydar, T.F., Kuan, C.Y., Flavell, R.A., and Rakic, P. (1999). The role of cell death in regulating the size and shape of the mammalian forebrain. Cereb. Cortex 9, 621–626.

Huang, H., Cotton, J.L., Wang, Y., Rajurkar, M., Zhu, L.J., Lewis, B.C., and Mao, J. (2013). Specific requirement of Gli transcription factors in Hedgehog-mediated intestinal development. J. Biol. Chem. 288, 17589–17596.

Huang, J., Wu, S., Barrera, J., Matthews, K., and Pan, D. (2005). The Hippo signaling pathway coordinately regulates cell proliferation and apoptosis by inactivating Yorkie, the Drosophila Homolog of YAP. Cell 122, 421–434.

Huangfu, D., and Anderson, K.V. (2005). Cilia and Hedgehog responsiveness in the mouse. Proceedings of the National Academy of Sciences 102, 11325–11330.

Huycke, T.R., Miller, B.M., Gill, H.K., Nerurkar, N.L., Sprinzak, D., Mahadevan, L., and Tabin, C.J. (2019). Genetic and Mechanical Regulation of Intestinal Smooth Muscle Development. Cell 179, 90–105.e21.

Ishizuya-Oka, A., and Ueda, S. (1996). Apoptosis and cell proliferation in the Xenopus small intestine during metamorphosis. Cell Tissue Res. 286, 467–476.

Jacoby, M., Cox, J.J., Gayral, S., Hampshire, D.J., Ayub, M., Blockmans, M., Pernot, E., Kisseleva, M.V., Compère, P., Schiffmann, S.N., et al. (2009). INPP5E mutations cause primary cilium signaling defects, ciliary instability and ciliopathies in human and mouse. Nat. Genet. 41, 1027–1031.

Kablar, B., Tajbakhsh, S., and Rudnicki, M.A. (2000). Transdifferentiation of esophageal smooth to skeletal muscle is myogenic bHLH factor-dependent. Development 127, 1627–1639.

Khalipina, D., Kaga, Y., Dacher, N., and Chevalier, N.R. (2019). Smooth muscle contractility causes the gut to grow anisotropically. Journal of the Royal Society Interface 16, 20190484.

Kim, H.Y., Pang, M.-F., Varner, V.D., Kojima, L., Miller, E., Radisky, D.C., and Nelson, C.M. (2015). Localized Smooth Muscle Differentiation Is Essential for Epithelial Bifurcation during Branching Morphogenesis of the Mammalian Lung. Developmental Cell 34, 719–726.

Kobelev, A.V., Smoluk, L.T., Lookin, O.N., Balakin, A.A., and Protsenko, Y.L. (2011). Modeling of steady-state and relaxation elastic properties of the papillary muscle at rest. Biophysics 56, 502–509.

Komarova, I.A., and Vorob’ev, I.A. (1995). [The centrosome structure in enterocytes in the histogenesis of the mouse intestine]. Ontogenez 26, 390–399.

Kuida, K., Haydar, T.F., Kuan, C.Y., Gu, Y., Taya, C., Karasuyama, H., Su, M.S., Rakic, P., and Flavell, R.A. (1998). Reduced apoptosis and cytochrome c-mediated caspase activation in mice lacking caspase 9. Cell 94, 325–337.

Lahiry, P., Wang, J., Robinson, J.F., Turowec, J.P., Litchfield, D.W., Lanktree, M.B., Gloor, G.B., Puffenberger, E.G., Strauss, K.A., Martens, M.B., et al. (2009). A Multiplex Human Syndrome Implicates a Key Role for Intestinal Cell Kinase in Development of Central Nervous, Skeletal, and Endocrine Systems. The American Journal of Human Genetics 84, 822.

le Duc, Q., Shi, Q., Blonk, I., Sonnenberg, A., Wang, N., Leckband, D., and de Rooij, J. (2010). Vinculin potentiates E-cadherin mechanosensing and is recruited to actin-anchored sites within adherens junctions in a myosin II-dependent manner. J. Cell Biol. 189, 1107–1115.

Lei, Q.-Y., Zhang, H., Zhao, B., Zha, Z.-Y., Bai, F., Pei, X.-H., Zhao, S., Xiong, Y., and Guan, K.-L. (2008). TAZ promotes cell proliferation and epithelial-mesenchymal transition and is inhibited by the hippo pathway. Mol. Cell. Biol. 28, 2426–2436.

Liem, K.F., Ashe, A., He, M., Satir, P., Moran, J., Beier, D., Wicking, C., and Anderson, K.V. (2012). The IFT-A complex regulates Shh signaling through cilia structure and membrane protein trafficking. J. Cell Biol. 197, 789–800.

Liu, A., Wang, B., and Niswander, L.A. (2005). Mouse intraflagellar transport proteins regulate both the activator and repressor functions of Gli transcription factors. Development 132, 3103–3111.

Lopez-Rios, J., Speziale, D., Robay, D., Scotti, M., Osterwalder, M., Nusspaumer, G., Galli, A., Holländer, G.A., Kmita, M., and Zeller, R. (2012). GLI3 constrains digit number by controlling both progenitor proliferation and BMP-dependent exit to chondrogenesis. Developmental Cell 22, 837–848.

Madison, B.B., Braunstein, K., Kuizon, E., Portman, K., Qiao, X.T., and Gumucio, D.L. (2005). Epithelial hedgehog signals pattern the intestinal crypt-villus axis. Development 132, 279–289.

Mao, J., Kim, B.-M., Rajurkar, M., Shivdasani, R.A., and McMahon, A.P. (2010). Hedgehog signaling controls mesenchymal growth in the developing mammalian digestive tract. Development 137, 1721–1729.

Moon, H., Song, J., Shin, J.-O., Lee, H., Kim, H.-K., Eggenschwiller, J.T., Bok, J., and Ko, H.W. (2014). Intestinal cell kinase, a protein associated with endocrine-cerebro-osteodysplasia syndrome, is a key regulator of cilia length and Hedgehog signaling. Proc. Natl. Acad. Sci. U.S.a. 111, 8541–8546.

Motoyama, J., Heng, H., Crackower, M.A., Takabatake, T., Takeshima, K., Tsui, L.C., and Hui, C. (1998). Overlapping and non-overlapping Ptch2 expression with Shh during mouse embryogenesis. Mechanisms of Development 78, 81–84.

Oud, M.M., Bonnard, C., Mans, D.A., Altunoglu, U., Tohari, S., Ng, A.Y.J., Eskin, A., Lee, H., Rupar, C.A., de Wagenaar, N.P., et al. (2016). A novel ICK mutation causes ciliary disruption and lethal endocrine-cerebro-osteodysplasia syndrome. Cilia 5, 8.

Paige Taylor, S., Kunova Bosakova, M., Varecha, M., Balek, L., Barta, T., Trantirek, L., Jelinkova, I., Duran, I., Vesela, I., Forlenza, K.N., et al. (2016). An inactivating mutation in intestinal cell kinase, ICK, impairs hedgehog signalling and causes short rib-polydactyly syndrome. Hum. Mol. Genet. 25, 3998–4011.

Panciera, T., Azzolin, L., Cordenonsi, M., and Piccolo, S. (2017). Mechanobiology of YAP and TAZ in physiology and disease. Nat. Rev. Mol. Cell Biol. 18, 758–770.

Pazour, G.J., Dickert, B.L., Vucica, Y., Seeley, E.S., Rosenbaum, J.L., Witman, G.B., and Cole, D.G. (2000). Chlamydomonas IFT88 and Its Mouse Homologue, Polycystic Kidney Disease Gene Tg737, Are Required for Assembly of Cilia and Flagella. J. Cell Biol. 151, 709–718.

Porteous, S., Torban, E., Cho, N.P., molecular, H.C.H., 2000 (2000). Primary renal hypoplasiain humans and mice with PAX2 mutations: evidence of increased apoptosis in fetal kidneys of Pax21Neu +/− mutant mice. Hum. Mol. Genet. 9.

Raleigh, D.R., Choksi, P.K., Krup, A.L., Mayer, W., Santos, N., and Reiter, J.F. (2018). Hedgehog signaling drives medulloblastoma growth via CDK6. J. Clin. Invest. 128, 120–124.

Ramalho-Santos, M., Melton, D.A., and McMahon, A.P. (2000). Hedgehog signals regulate multiple aspects of gastrointestinal development. Development 127, 2763–2772.

Salbreux, G., Charras, G., and Paluch, E. (2012). Actin cortex mechanics and cellular morphogenesis. Trends Cell Biol. 22, 536–545.

Savin, T., Kurpios, N.A., Shyer, A.E., Florescu, P., Liang, H., Mahadevan, L., and Tabin, C.J. (2011). On the growth and form of the gut. Nature 476, 57–62.

Segel, M., Neumann, B.X.R., Hill, M.F.E., Weber, I.P., Viscomi, C., Zhao, C., Young, A., Agley, C.C., Thompson, A.J., Gonzalez, G.A., et al. (2019). Niche stiffness underlies the ageing of central nervous system progenitor cells. Nature 573, 1–27.

Shyer, A.E., Tallinen, T., Nerurkar, N.L., Wei, Z., Gil, E.S., Kaplan, D.L., Tabin, C.J., and Mahadevan, L. (2013). Villification: how the gut gets its villi. Science 342, 212–218.

Stark, R., and Dunn, J.C.Y. (2012). Mechanical Enterogenesis - A Review. Journal of Healthcare Engineering 3, 229–242.

Stooke-Vaughan, G.A., and Campàs, O. (2018). Physical control of tissue morphogenesis across scales. Curr. Opin. Genet. Dev. 51, 111–119.

Tong, Y., Park, S.H., Wu, D., Xu, W., Guillot, S.J., Jin, L., Li, X., Wang, Y., Lin, C.-S., and Fu, Z. (2017). An essential role of intestinal cell kinase in lung development is linked to the perinatal lethality of human ECO syndrome. FEBS Lett. 591, 1247–1257.

Tuveson, D., and Clevers, H. (2019). Cancer modeling meets human organoid technology. Science 364, 952–955.

Vichas, A., and Zallen, J.A. (2011). Translating cell polarity into tissue elongation. Semin. Cell Dev. Biol. 22, 858–864.

Vico, L., Collet, P., Guignandon, A., Lafage-Proust, M.H., Thomas, T., Rehaillia, M., and Alexandre, C. (2000). Effects of long-term microgravity exposure on cancellous and cortical weight-bearing bones of cosmonauts. Lancet 355, 1607–1611.

Walton, K.D., Freddo, A.M., Wang, S., and Gumucio, D.L. (2016). Generation of intestinal surface: an absorbing tale. Development 143, 2261–2272.

Wang, B., Zhang, Y., Dong, H., Gong, S., Wei, B., Luo, M., Wang, H., Wu, X., Liu, W., Xu, X., et al. (2018). Loss of Tctn3 causes neuronal apoptosis and neural tube defects in mice. Cell Death Dis 9, 520.

Wells, J.M., and Spence, J.R. (2014). How to make an intestine. Development 141, 752–760.

Wijgerde, M., McMahon, J.A., Rule, M., and McMahon, A.P. (2002). A direct requirement for Hedgehog signaling for normal specification of all ventral progenitor domains in the presumptive mammalian spinal cord. Genes & Development 16, 2849–2864.

Yang, Y., Roine, N., and Mäkelä, T.P. (2013). CCRK depletion inhibits glioblastoma cell proliferation in a cilium-dependent manner. EMBO Rep. 14, 741–747.

Yu, K., Xu, J., Liu, Z., Sosic, D., Shao, J., Olson, E.N., Towler, D.A., and Ornitz, D.M. (2003). Conditional inactivation of FGF receptor 2 reveals an essential role for FGF signaling in the regulation of osteoblast function and bone growth. Development 130, 3063–3074.

Zhao, B., Wei, X., Li, W., Udan, R.S., Yang, Q., Kim, J., Xie, J., Ikenoue, T., Yu, J., Li, L., et al. (2007). Inactivation of YAP oncoprotein by the Hippo pathway is involved in cell contact inhibition and tissue growth control. Genes & Development 21, 2747–2761.

Zhao, B., Ye, X., Yu, J., Li, L., Li, W., Li, S., Yu, J., Lin, J.D., Wang, C.-Y., Chinnaiyan, A.M., et al. (2008). TEAD mediates YAP-dependent gene induction and growth control. Genes & Development 22, 1962–1971.

Zhao, J.-B., Sha, H., Zhuang, F.-Y., and Gregersen, H. (2002). Morphological properties and residual strain along the small intestine in rats. World J. Gastroenterol. 8, 312–317.

Zheng, Y., and Pan, D. (2019). The Hippo Signaling Pathway in Development and Disease. Developmental Cell 50, 264–282.

